# Sortilin is a scavenger receptor for cytosolic proteins mislocalized to the extracellular milieu

**DOI:** 10.64898/2026.11.16.766851

**Authors:** Shotaro Sinchu, Kanna Kamikawa, Sayana Bun, Mako Kiyota, Akira Matsuura, Eisuke Itakura

**Affiliations:** Department of Biology, Graduate school of Science and Engineering, Chiba University, Chiba, Japan; Department of Biology, Graduate School of Science, Chiba University. Chiba, Japan; Center of Quantum Life Science for Structural Therapeutics (cQUEST), Chiba University, Chiba, Japan; Institute for Advanced Academic Research (IAAR), Chiba University, Chiba, Japan

**Keywords:** Protein quality control, sortilin, lysosomal degradation, hetero-autophagy, glycosylated protein

## Abstract

Aberrant intracellular proteins triggered by various abnormalities, such as misfolding or mislocalization, are degraded by protein quality-control (PQC) systems. However, how extracellular PQC (ePQC) recognizes and eliminates cytosolic proteins —leaked and extracellularly mislocalized upon tissue injury—remains poorly understood. Here, we identify sortilin as a novel ePQC scavenger receptor that preferentially recognizes leaked, non-glycosylated cytosolic proteins in the extracellular space as abnormal and targets them for lysosomal degradation. Through hydrophobic interactions mediated by multiple hydrophobic patches on the sortilin surfaces, sortilin broadly engages diverse cytosolic substrates (e.g., hemoglobin), while discriminating against secreted proteins protected by hydrophilic glycans. Crucially, this clearance is driven by a “hit-and-run” shuttling mechanism: sortilin resides at the cell surface for only a few minutes, and this rapid cycling, combined with broad-spectrum recognition, enables the degradation of even low-affinity substrates. Together, these findings establish sortilin as an ePQC scavenger receptor for extracellularly mislocalized cytosolic proteins.

## Introduction

To maintain homeostasis, a protein quality-control (PQC) system must effectively recognize and degrade a diverse range of abnormal proteins, including misfolded, aberrantly modified, or inappropriately localized species. Proteins with these distinct defects are targeted by specific PQC pathways. For example, molecular chaperones identify misfolded proteins and direct them toward either refolding or degradation^1^.

Within the endoplasmic reticulum (ER), the accumulation of immature proteins (e.g., those that are incompletely glycosylated or lack disulfide bonds) triggers the unfolded protein response (UPR), leading to their elimination via ER-associated degradation (ERAD)^2,3^. Mislocalized organelle proteins are specifically recognized and targeted for rapid degradation; for example, ER and mitochondrial proteins are promptly cleared upon mistargeting to non-cognate organelles^4–11^. When invasive bacteria rupture the endosome to enter the cytoplasm, the exposed glycans of endosomal membrane proteins serve as danger signals recognized by cytoplasmic galectins, which then trigger autophagic bacterial degradation^12,13^. Together, the major intracellular proteolytic systems—the ubiquitin-proteasome machinery and the autophagy-lysosome pathway—maintain robust intracellular PQC by specifically recognizing and degrading aberrant proteins. In addition, autophagy constantly turns over cytosolic proteins through non-selective degradation, thereby enabling the stochastic degradation of the entire cytosolic protein pool^14^. Dysfunction of these systems is implicated in a broad spectrum of human disorders, including neurodegenerative diseases, cancer, immunological conditions, and metabolic diseases^1,15^.

Protein levels are high not only intracellularly but also extracellularly, notably in body fluids such as blood. Indeed, about 11% of human genes encode secreted proteins^16^, many of which play essential physiological roles. As in the intracellular environment, extracellular accumulation of aberrant proteins is a critical hallmark of disease. For instance, Alzheimer’s disease, which affects about 32 million people worldwide^17^, is primarily characterized by the accumulation of extracellular amyloid-beta plaques. Our recent work on extracellular protein quality control (ePQC) for misfolded proteins revealed a pathway by which scavenger carriers—clusterin, α2-macroglobulin, and GPLD1— recognize misfolded extracellular proteins and direct them for lysosomal degradation via endocytosis^18–20^. However, abnormal extracellular proteins are not restricted to misfolded species. Intracellular components are routinely leaked into the extracellular space under physiological conditions, such as basal hemolysis^21^, and are heavily released during tissue disruption accompanied by necrosis, pyroptosis, or plasma membrane rupture in pathological states, including myocardial infarction, acute liver injury, rhabdomyolysis, and ischemia-reperfusion injury^22–25^. While a specific subset of these leaked cytosolic proteins acts as damage-associated molecular patterns (DAMPs) that engage receptors such as RAGE to activate macrophages and immune responses^26^, most mislocalized cytosolic proteins in the extracellular space remain natively folded. Whether and how such a vast repertoire of mislocalized cytosolic proteins is comprehensively recognized and cleared from the extracellular milieu remains virtually unknown.

We previously used a comprehensive CRISPR knockout (KO) genetic screen to identify genes required for the internalization of clusterin-RFP-GFP and found that heparan sulfate, a glycosaminoglycan, acts as a receptor of the clusterin–substrate complex^18^. Re-evaluation of this dataset identified SORT1 (encoding sortilin) as another robust candidate (data not shown). Sortilin is a single-transmembrane protein that serves as a multifunctional regulator of protein sorting to lysosomes and the plasma membrane^27^. Over 50 different proteins have been identified as sortilin-binding partners^28^. To test whether sortilin is a clusterin-specific receptor, we performed an endocytosis assay using clusterin-RFP-GFP in sortilin-overexpressing (OE) cells, employing RFP-GFP alone as a negative control. Surprisingly, uptake of the RFP-GFP control was markedly enhanced in sortilin-OE cells. This led us to hypothesize that sortilin functions as a broad-spectrum receptor targeting a wide range of proteins in the extracellular space.

In the present study, we elucidate how mislocalized cytosolic proteins are recognized and cleared in the extracellular space. Although most secreted proteins are glycosylated, we found that extracellular non-glycosylated proteins, including leaked cytosolic proteins, are directly recognized by cell-surface sortilin and then preferentially eliminated via lysosomal degradation. We show that sortilin utilizes surface hydrophobic patches (HPs) for broad substrate recognition and undergoes rapid membrane-to-endosome cycling, operating as a “hit-and-run” shuttle that captures and delivers a broad spectrum of extracellular proteins to lysosomes for degradation. Our results highlight a novel ePQC that facilitates the degradation of non-glycosylated proteins recognized as mislocalized cytosolic proteins in the extracellular milieu.

## Results

### Sortilin facilitates extensive protein internalization

To investigate the lysosomal degradation of extracellular proteins, we established a system to monitor their trafficking to the lysosome. As illustrated in Figure 1A, this system utilizes a tandem fluorescent protein (RFP-GFP), in which mCherry (RFP) is resistant to lysosomal proteases and sfGFP (GFP) is sensitive^29^. Upon endocytosis and subsequent transport to the lysosome, the GFP moiety is degraded, whereas the stable RFP and the ATTO-488 and −568 fluorophores accumulate, indicating successful lysosomal delivery (Figure 1A).

**Figure 1.**
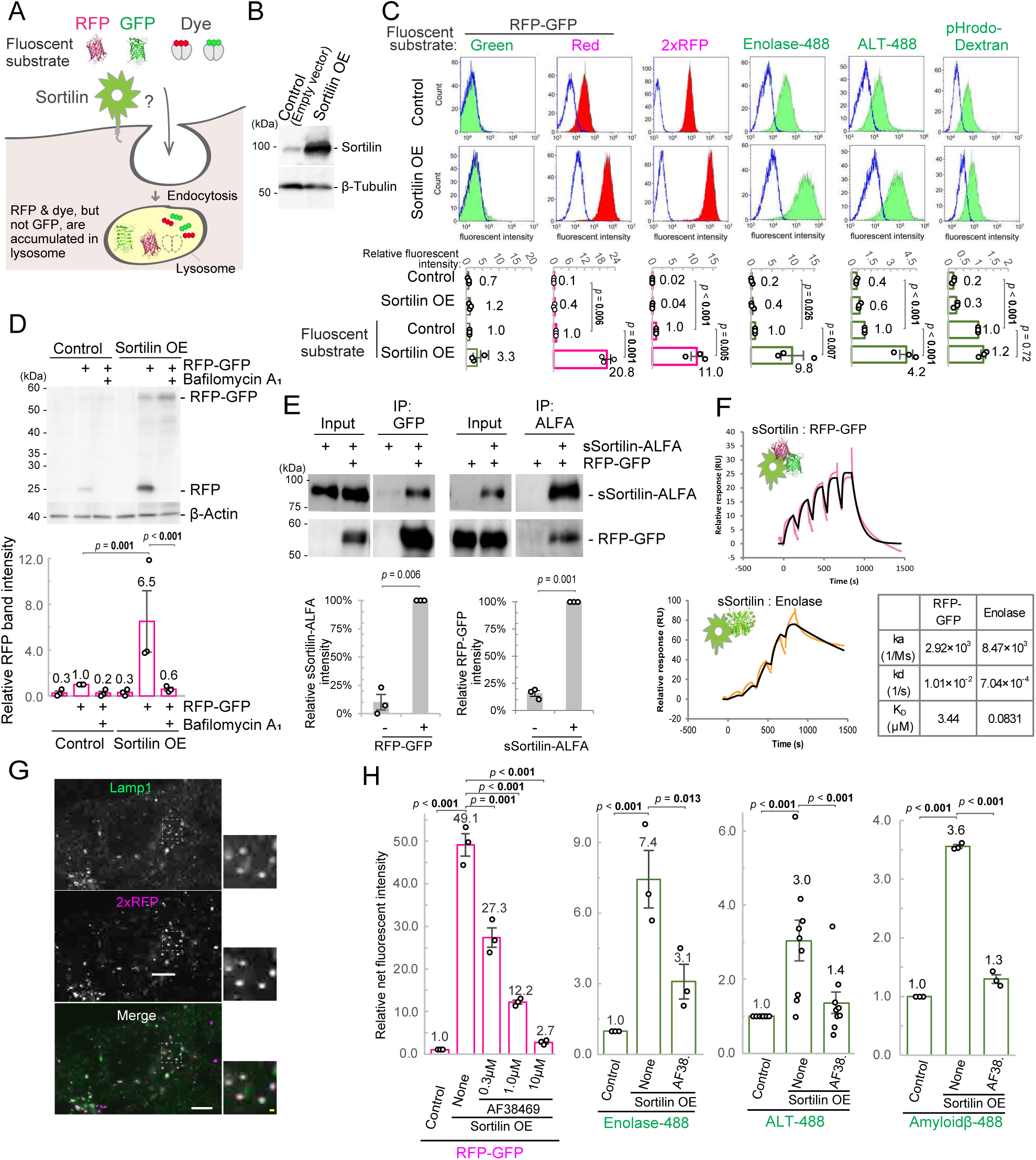
Sortilin facilitates lysosomal degradation of extracellular proteins. (A) Schematic of the endocytosis assay. After internalization, lysosomal enzymes degrade GFP, whereas RFP and the dye, which are resistant to proteases and acidic pH, accumulate in lysosomes. (B) Immunoblot of control and Sortilin-overexpressing (OE) (HEK293) cells using anti-sortilin and -β-tubulin antibodies. (C) Flow cytometry (FCM) analysis of fluorescent extracellular protein endocytosis in control and sortilin OE cells (HEK293) incubated with the indicated proteins for 16 h. Fluorescent intensity per cell as measured by FCM. The blue dashed line is the signal from untreated control cells. This defines the extent of cellular autofluorescence. (D) Immunoblot analysis of RFP cleavage in control and sortilin OE cells (HEK293). Control and sortilin OE cells (HEK293) were incubated with 0.5 μM RFP-GFP in the presence or absence of bafilomycin A1 for 16 h. Relative RFP intensity is shown as the fold change relative to RFP-GFP-treated control cells, which was set as 1. (E) In vitro immunoprecipitation of sSortilin and RFP-GFP. sSortilin-ALFA-His and RFP-GFP were mixed and subjected to immunoprecipitation using anti-GFP nanobody beads. The resulting precipitates were then subjected to immunoblotting using antibodies against GFP and ALFA. The relative band intensity is the -fold change of each sample intensity, normalized to the intensity of the sample containing both proteins, which was set to 100%. (F) SPR sensorgrams showing binding kinetics of sSortilin to RFP-GFP and enolase. Substrates (RFP-GFP or enolase) were sequentially injected over captured sSortilin using a single-cycle kinetics protocol. Experimental data are represented by colored lines with fitted curves depicted in black, derived from a 1:1 binding model. (G) Confocal images of internalized 2xRFP and a lysosome marker (Lamp1). Sortilin OE cells (HeLa) were incubated with 0.5 μM 2xRFP for 16 h. (H) Inhibition of sortilin-dependent internalization by a sortilin inhibitor, AF38469 (AF38.). Control or sortilin OE cells (HEK293) were incubated with the indicated proteins in the presence or absence of 0.3 - 10 μM (for RFP-GFP) or 10 μM AF38469 (for enolase, ALT, Amyloid-β) for 16 h and analyzed by FCM. The relative (net) fluorescent intensity is the -fold change of the net fluorescent intensity of each sample normalized to the (net) fluorescent intensity of control cells, which was set to 1. Data are presented as means ± s.e.ms.; n = 3 [C, D, E, H (for RFP-GFP, enolase, Amyloid-β)], n = 9 [H (for ALT)]. Scale bars: 10 μm, 2 μm (inset).

We hypothesized that sortilin is a versatile receptor, recognizing a broad array of proteins or serving as a general activator of endocytosis. To test this hypothesis, we generated stable sortilin-overexpressing (OE) cells, which expressed sortilin at levels more than 10-fold higher than those of endogenous sortilin in control cells (with an empty vector), as confirmed by immunoblotting (Figure 1B). To evaluate the direct endocytic uptake of extracellular proteins, cells were incubated with fluorescently labeled substrates (0.01–0.5 µM) in serum-free medium supplemented with minimal growth factors (albumin, insulin, and transferrin) to prevent starvation without the influence of uncharacterized serum.

Flow cytometry (FCM) analysis revealed a selective increase in the RFP signal in RFP-GFP–treated cells, confirming lysosomal delivery (Figure 1C). We also observed significant increases in the uptake of several proteins, including 2xRFP, albumin-565, and amyloid-β-488 (Figures 1C and S1A). Even enolase, alanine aminotransferase (ALT/GPT), and γ-glutamyl transferase (γ-GT), which serve as clinical serum markers of tissue damage^30–32^, also underwent sortilin-dependent endocytosis. Notably, the uptake of dextran (a fluid-phase marker) was not significantly increased, suggesting that sortilin OE enhanced the internalization of a broad range of proteins rather than activating general endocytosis. In contrast, the increase in transferrin internalization in sortilin OE cells was only marginal (Figure S1A), implying that some proteins do not depend on sortilin for internalization.

In addition, the minor increase in autofluorescence in sortilin-OE cells compared with wild-type (WT) cells without exogenous substrates (blue dashed line in Figure 1C and S1A) suggests the accumulation of degradation intermediates in lysosomes. Notably, even with our previously developed lysosomal-METRIQ assay^33^, sortilin OE showed no detrimental effect on lysosomal activity (Figure S1B), indicating that the increased autofluorescence reflects an increase in intralysosomal degradation intermediates driven by an elevated flux of internalized proteins, rather than lysosomal dysfunction.

We next examined the uptake and accumulation of other fluorescent proteins in sortilin-OE cells, including 1xGamillus (a pH-and protease-resistant GFP)^34^, 2xGamillus, 1xRFP, and 2xRFP (Figure S1C). While the internalization of 1xGamillus or 1xRFP by sortilin-OE cells was 4.6 and 2.5-fold higher than in control cells, the uptake of 2xGamillus and 2xRFP was increased more than 20 and 11-fold, respectively. We speculate that this is attributable to the increased avidity caused by dimerization.

Degradation of internalized 2xFP (56 kDa; 2x fluorescent protein) was biochemically confirmed by the appearance of a 25-kDa RFP monomer, the size expected for cleaved RFP, in lysosomes (RFP cleavage assay). Cleaved RFP was increased in sortilin OE cells, and this increase was completely inhibited by the lysosomal inhibitor bafilomycin A1, a V-ATPase inhibitor that blocks endocytic and lysosomal pathways^35,36^ (Figure 1D). These data indicate that the effects of sortilin overexpression are due to enhanced internalization of extracellular proteins, rather than the inhibition of lysosomal enzymes. Importantly, fluorescent proteins lack a physiological receptor on mammalian cells, and their delivery to the lysosome can be biochemically detected via the RFP cleavage assay. Thus, 2xFP serves as an ideal marker for monitoring both sortilin-dependent endocytosis and subsequent lysosomal degradation.

*In vitro* immunoprecipitation using purified soluble sortilin (sSortilin), which comprises the entire extracellular domain (amino acids 1–753), and RFP-GFP showed direct binding between sSortilin-ALFA and RFP-GFP (Figure 1E). Surface plasmon resonance (SPR) binding kinetics revealed that sortilin exhibits a wide affinity range for its substrates, from moderate-to-high affinity for enolase (KD = 0.08 µM) to weak binding for RFP-GFP (KD = 3.44 µM) (Figure 1F). To confirm the destination of internalized proteins, immunofluorescence staining showed that internalized 2xRFP co-localized with the lysosomal marker Lamp1 (Figure 1G). We also investigated whether AF38469, a sortilin inhibitor that targets the neurotensin binding site^37^, affects protein internalization in sortilin-OE cells. AF38469 treatment suppressed the endocytosis of RFP-GFP in a concentration-dependent manner (Figure 1H). Furthermore, the uptake of other fluorescently labeled proteins—enolase-488, ALT-488, and amyloid-β-488—was substantially blocked by AF38469 in sortilin-OE cells. Taken together, these data indicate that sortilin facilitates the lysosomal degradation of a broad array of proteins in the extracellular space.

### Sortilin and SorLA drive protein internalization

To explore whether other members of the sortilin family (SorLA, SorCS1, SorCS2, SorCS3)^27,38–42^ were involved in this process (Figure 2A), we generated cell lines stably overexpressing each sortilin family member (Figure 2B) and performed endocytosis assays. Overexpression of sortilin and SorLA significantly increased the uptake of exogenously added proteins, while that of SorCS1–3 exerted only minor effects (Figures 2C and S2A).

**Figure 2.**
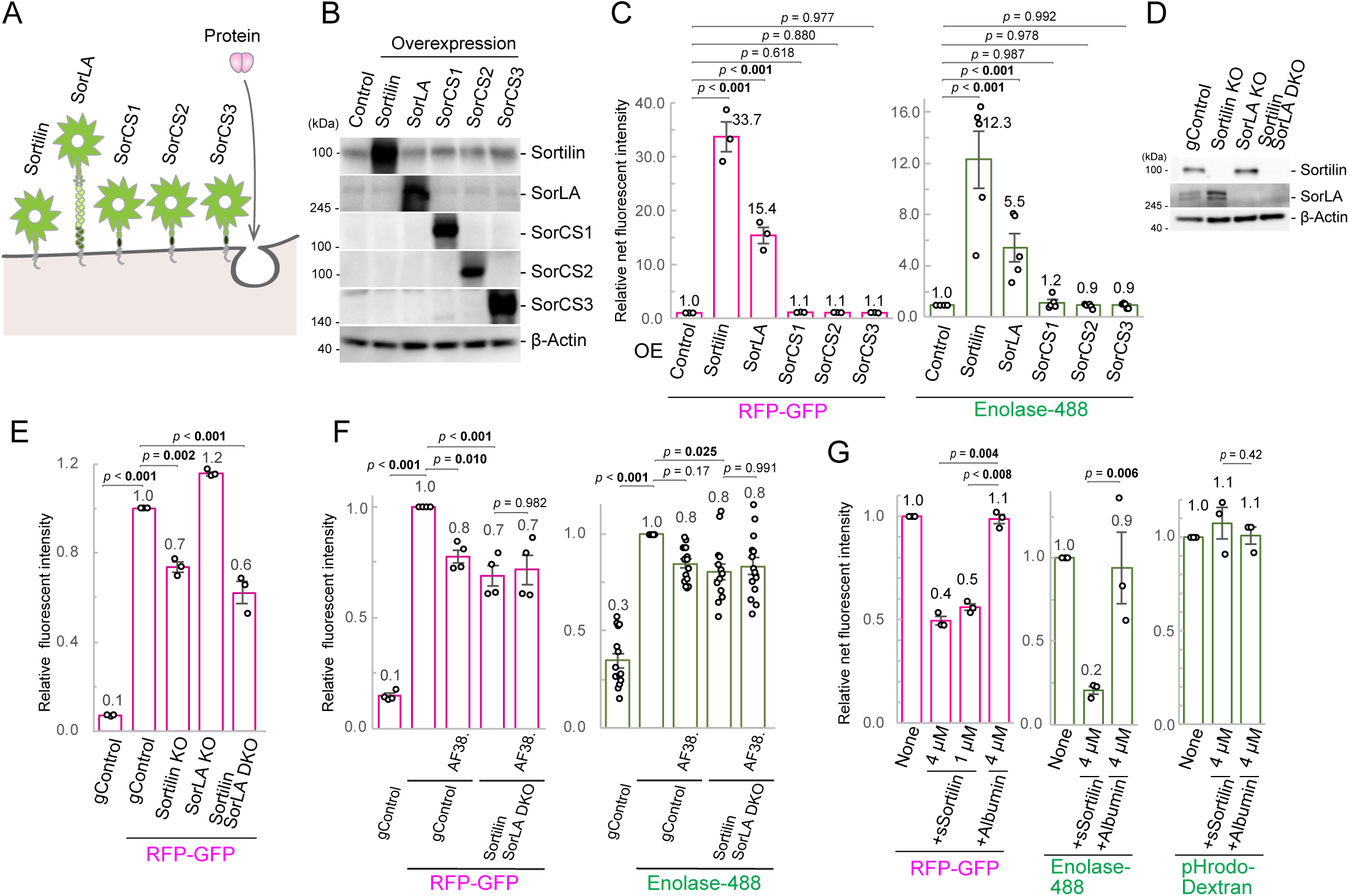
Sortilin and SorLA are required for internalization of extracellular proteins. (A) Schematic representation of the endocytosis assay using cells expressing sortilin, SorLA, SorCS1, SorCS2, or SorCS3. (B) Immunoblot of control and sortilin family-overexpressing (OE) cells (HEK293) using the indicated antibodies. (C) Fluorescent extracellular protein internalization by control and sortilin family OE cells (HEK293) incubated with the indicated proteins for 16 h, and analyzed by FCM. (D) Immunoblot of control guide RNA (gControl), sortilin KO, SorLA KO, and sortilin and SorLA DKO (HEK293) cells using indicated antibodies. (E) Fluorescent extracellular protein internalization in gControl and KO cells incubated with 0.5 μM RFP-GFP for 16 h, and analyzed by FCM. (F) Control and sortilin and SorLA DKO cells (HEK293) incubated with the indicated proteins in the presence or absence of 10 μM AF38469 for 16 h, and analyzed by FCM. (G) Competitive inhibition of fluorescent protein internalization by sSortilin. HEK293 cells were incubated with 0.2 μM RFP-GFP, 0.01 μM enolase-488 or 0.2 ng/ul pHrodo-Dextran in the presence or absence of sSortilin-ALFA or albumin for 16 h and analyzed by FCM. The relative (net) fluorescent intensity is the -fold change of the (net) fluorescent intensity of each sample normalized to the (net) fluorescent intensity of control cells (C), gControl cells (E and F) or none (G), all of which were set to 1. Data are presented as means ± s.e.ms.; n = 3 [C (for RFP-GFP), E, G], n = 4 [F (for RFP-GFP)], n = 5 [C (for enolase)], n = 15 [F (for enolase)].

To confirm the physiological roles of sortilin and SorLA, we generated single KO and double KO (DKO) cell lines using CRISPR/Cas9 (Figure 2D). Endocytosis assays using these cells revealed that endogenous sortilin mediated extracellular protein uptake, whereas the contribution of endogenous SorLA was minimal (Figures 2E and S2B). Uptake of enolase was also significantly decreased in DKO cells (Figure 2F). Crucially, the reduction in uptake induced by the sortilin inhibitor AF38469 was abolished in DKO cells, verifying the target specificity of AF38469 for sortilin (Figures 2F and S2C). In addition, the impaired endocytosis in DKO cells was rescued by the re-expression of either sortilin or SorLA (Figure S2D). The residual substrate uptake observed even in DKO cells is likely attributable to alternative uptake mechanisms, such as fluid-phase endocytosis.

Sortilin acts not only as a cell-surface endocytic receptor but also as an intracellular sorting receptor at the Golgi apparatus^27^. To selectively inhibit cell-surface sortilin, we added excess (1 or 4 μM) sSortilin to the culture medium. This competitively inhibited the internalization of RFP-GFP, 2xGamillus, 2xRFP, albumin-565, amyloid-β-488, enolase-488, and ALT-488, but not transferrin-488 or dextran (Figures 2G and S2E), suggesting that sortilin serves as a cell-surface receptor for a wide variety of proteins.

We next examined the expression levels of endogenous sortilin, SorLA, and SorCS1–3 in various cell lines. Sortilin was broadly expressed (Figure S3A), whereas SorLA and SorCS2 were detected in only a limited number of cell lines. SorCS1 and SorCS3 were undetectable in all cells examined. In addition, sortilin was barely detectable in macrophage cell lines (Figure S3B). Although all cell lines tested exhibited internalization, the correlation between sortilin expression and uptake rate differed markedly among substrates (Figure S3C). In particular, sortilin expression showed only a modest correlation with the uptake of fluorescent proteins (RFP-GFP and 2xRFP), whereas virtually no correlation was observed for enolase-488, ALT-488, or amyloid-β-488. This lack of correlation suggests the contribution of auxiliary factors, such as substrate-specific receptors and distinct basal endocytic rates. While RFP-GFP and 2xRFP may not offer absolute sortilin specificity, they remain better markers than the other substrates for evaluating sortilin-dependent uptake and degradation.

Given the high levels of albumin (∼500–750 µM) in blood^43^ and our data showing that albumin undergoes sortilin-dependent endocytosis (Figures S2A and S2E), we investigated whether high albumin concentrations affect the sortilin-dependent endocytosis of other substrates. The sortilin-dependent uptake of 0.5 µM RFP-GFP, 0.01 µM enolase-488, and 0.01 µM ALT-488 was largely unaffected, showing only a minimal decrease, even in the presence of 5–500 µM albumin (Figure S3D). These findings demonstrate that albumin barely competes with these substrates for sortilin-dependent endocytosis.

### Sortilin targets non-glycosylated proteins

These results raised fundamental questions: Why is the degradation of fluorescent protein facilitated by sortilin? What defines a protein as “abnormal” in the extracellular space? Most extracellular proteins are glycosylated, whereas our reporter proteins were not. We therefore hypothesized that sortilin facilitates the degradation of proteins lacking glycans. To test this, we used IgE, which has 12 glycosylation sites, and IgG, which has only two^44^. SDS-PAGE confirmed that PNGase F treatment, which removes N-glycans, caused a significant upward shift in IgE mobility but had a minor effect on IgG (Figure 3A). FCM showed that IgE uptake was significantly increased after PNGase F treatment compared with that of IgG, and this enhanced uptake was competitively inhibited by excess extracellular sSortilin (Figure 3B). In addition, sortilin OE significantly increased the uptake of PNGase F-treated IgE (Figure 3C), and immunofluorescence staining confirmed that the PNGase F-treated IgE was delivered to the lysosomes (Figure 3D).

**Figure 3.**
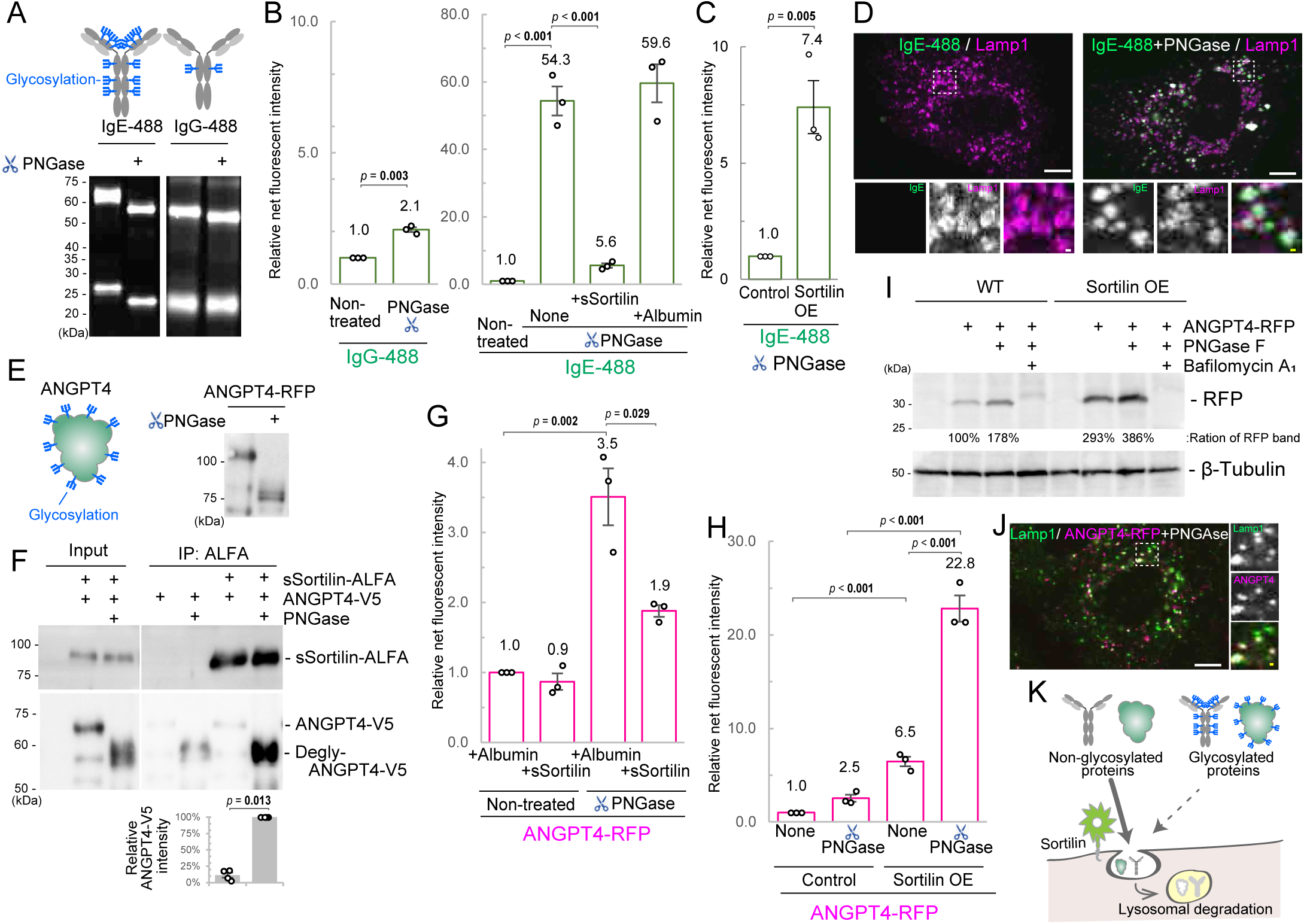
Non-glycosylated proteins are preferentially internalized by sortilin. (A) IgE has 12 glycosylation sites and IgG has two. IgE-488 and IgG-488 were pre-treated with PNGase F for 3 h at 37°C and then subjected to SDS-PAGE and fluorescence imaging. (B) FCM analysis of deglycosylated IgG-488 and IgE-488 internalization. Human umbilical vein endothelial cells (HuEht-2 line) were incubated with 2 ng/µL IgG-488 or 0.3 ng/µL IgE-488 with or without 2 μM sSortilin-ALFA or 2 μM albumin for 16 h, and analyzed by FCM. (C) FCM analysis of IgE in sortilin OE cells. Control and Sortilin OE cells (HEK293) were incubated with 0.3 ng/µL IgE-488 (PNGase F pre-treated) for 16 h, and analyzed by FCM. (D) Confocal images of internalized IgE-488 and a lysosome marker (Lamp1). HuEht-2 cells were incubated with (PNGase F-treated or -untreated) 0.3 ng/µL IgE-488 for 16 h. (E) ANGPT4 has nine glycosylation sites. ANGPT4-RFP was treated with PNGase F for 3 h at 37°C and subjected to immunoblotting using the anti-mCherry antibody. (F) In vitro immunoprecipitation of sSortilin with ANGPT4. sSortilin-ALFA-His, and glycosylated ANGPT4-V5 or deglycosylated ANGPT4-V5, were mixed and subjected to immunoprecipitation using anti-ALFA nanobody beads. The resulting precipitates were then immunoblotted using antibodies against V5, and ALFA. (G) Competitive inhibition of ANGPT4 internalization by sSortilin. HEK293 cells were incubated with 0.2 μM ANGPT4-RFP (PNGase F-treated or -untreated) in the presence or absence of 2 μM sSortilin-ALFA or 2 μM albumin for 16 h, and analyzed by FCM. (H) FCM analysis of sortilin-dependent internalization of deglycosylated ANGPT4. Control and sortilin OE cells (HEK293) were incubated with 0.2 μM ANGPT4-RFP (PNGase F-treated or -untreated) for 16 h, and analyzed by FCM. (I) Immunoblot analysis of the ANGPT4-RFP cleavage assay data on control and Sortilin OE cells (HEK293). Control and Sortilin OE cells (HEK293) were incubated with 0.2 μM ANGPT4-RFP with or without pre-treated PNGase F (for 3 h at 37°C) in the presence or absence of bafilomycin A1 for 16 h and immunoblotted using anti-mCherry and -β-tubulin antibodies. (J) Confocal images of internalized ANGPT4-RFP with a lysosome marker (Lamp1). Sortilin OE cells (HeLa) were incubated with PNGase F-treated 0.2 μM ANGPT4-RFP for 16 h. (K) A schematic model illustrating that sortilin mediates the endocytosis of non-glycosylated proteins, but less efficiently that of glycosylated proteins. The relative net fluorescent intensity in FCM is the -fold change of the net fluorescent intensity of each sample normalized to the net fluorescent intensities of non-treated cells (B), control cells (C), albumin-non-treated cells (G), or none in control cells (H), all of which were set to 1. Data are presented as means ± s.e.ms.; n = 3 (B, C, G, H), n = 4 (F). Scale bars: 10 μm, 2 μm (inset).

We next investigated ANGPT4, a secreted protein with nine glycosylation sites (Figure 3E). Recombinant ANGPT4-RFP was purified from the conditioned medium of mammalian cells. Notably, PNGase F treatment of ANGPT4 enhanced its binding to sSortilin (Figure 3F), suggesting that glycosylation interfered with sortilin binding. ANGPT4-RFP internalization was also enhanced by PNGase F treatment and competitively inhibited by sSortilin (Figure 3G). Uptake was also increased in sortilin-OE cells (Figure 3H). The RFP cleavage assay demonstrated that lysosomal degradation of ANGPT4-RFP was dependent on both PNGase F treatment and sortilin (Figure 3I). Co-localization with Lamp1 confirmed that internalized ANGPT4 was degraded in lysosomes (Figure 3J). In a control experiment, an identical PNGase F treatment did not enhance the internalization of RFP-GFP, which lacks glycosylation sites (Figure S4A), confirming that deglycosylation was required for this enhancement.

To determine whether ANGPT4-RFP internalization was mediated by a specific sugar moiety, we treated the protein with various glycosidases (neuraminidase A, galactosidase S, and acetylhexosaminidase f), which cleave distinct sugar moieties (Figures S4B and C)^45^. All three enzymes slightly increased ANGPT4-RFP internalization by sortilin-OE cells, but much less than the dramatic rise seen after PNGase F treatment (Figure S4D). This suggested that sortilin-dependent internalization was not mediated by recognition of a specific sugar residue but, rather, by the absence of N-glycans. Collectively, these data show that a lack of glycosylation is the key signal for cellular protein uptake by cell-surface sortilin (Figure 3K).

### Sortilin clears leaked cytosolic proteins mislocalized to the extracellular space

Taken together, our results indicate that non-glycosylated proteins are degraded by sortilin. The most abundant non-glycosylated proteins *in vivo* are likely of cytoplasmic origin, and enolase and ALT are cytosolic proteins that leak into the extracellular environment during tissue damage^30,32,46^. The aberrant release of cytosolic proteins into the extracellular space is a hallmark of various pathological conditions, including hemolytic disorders (e.g., sickle cell disease, hemolytic uremic syndrome, and malaria) and tissue injuries (e.g., rhabdomyolysis, acute liver failure, acute pancreatitis, and ischemia-reperfusion injury)^47,48^. To test whether sortilin promotes the endocytic degradation of cytosolic proteins leaked from tissues, we isolated cytosolic fractions from various mouse tissues (liver, muscle, and red blood cells (RBC)) (Figures S5) and labeled the crude cytosolic protein preparations with a fluorescent dye (Figure 4A). Endocytosis assays demonstrated that sortilin-OE cells exhibited significantly increased uptake of cytosolic proteins from all tissues tested (Figure 4B). This uptake was efficiently inhibited by the addition of excess sSortilin (Figure 4C).

**Figure 4.**
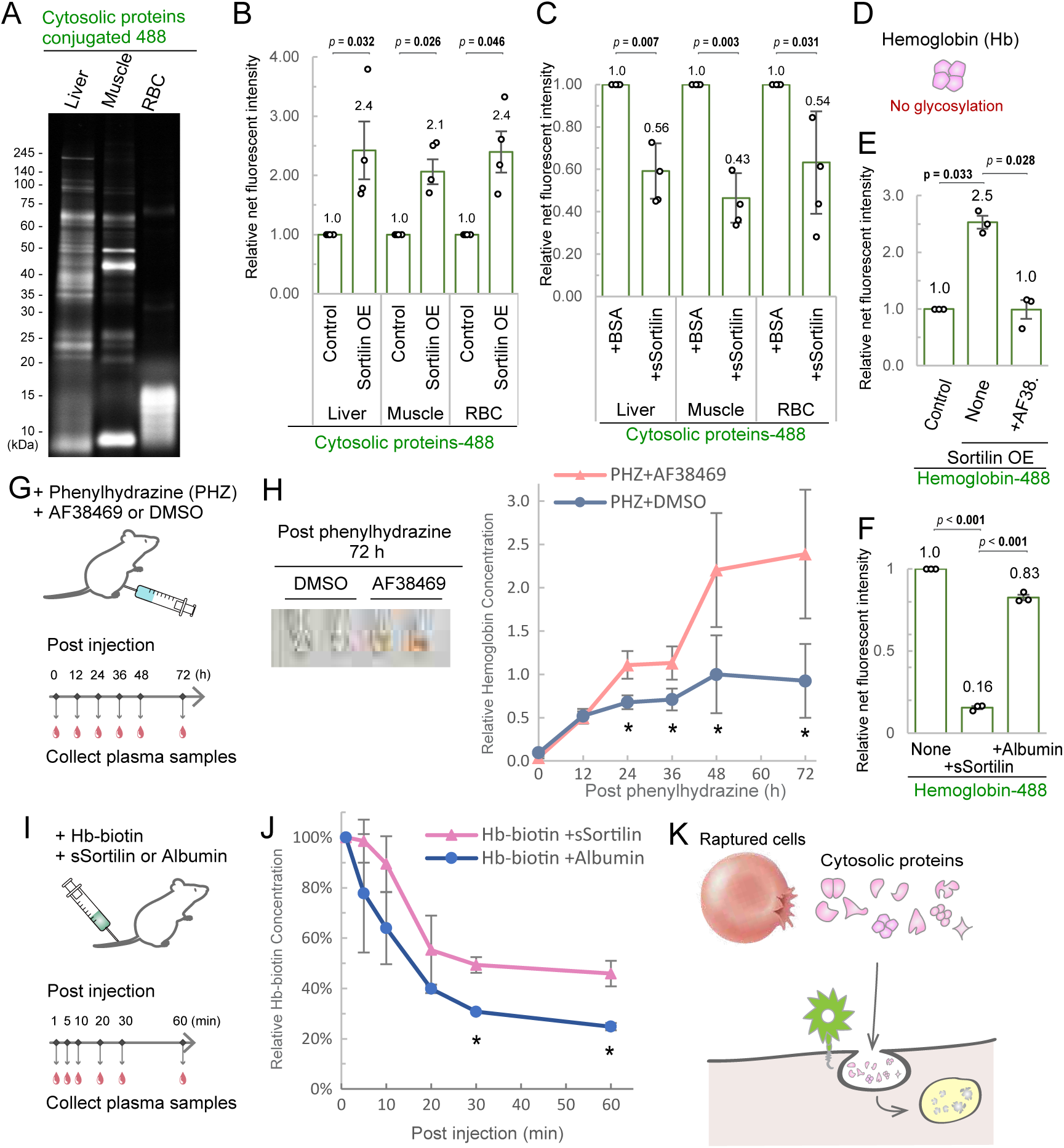
Clearance of leaked cytosolic proteins is mediated by sortilin in vitro and in vivo. (A) Fluorescent labeling of cytosolic proteins isolated from various tissues. The indicated tissue lysates were conjugated with NHS-ATTO 488 and subjected to SDS-PAGE and fluorescence imaging. (B) FCM analysis of internalization of cytosolic proteins from various tissues. Control and sortilin OE cells (HEK293) were incubated with 12 ng/µL of the indicated cytosolic proteins-488 for 16 h, and analyzed by FCM. (C) Competitive inhibition of cytosolic protein internalization by sSortilin. HEK293 cells were incubated with 12 ng/µL of the indicated cytosolic proteins-488 in the presence or absence of 2 μM sSortilin-ALFA or 2 μM albumin for 16 h, and analyzed by FCM. (D) Hb is a non-glycosylated protein. (E and F) FCM analysis of sortilin-dependent Hb internalization. Control and sortilin OE cells (HEK293) were incubated with 0.01 µM Hb-488 in the presence or absence of 10 μM AF38469 for 16 h, and analyzed by FCM. (E). HuEhT-2 cells were incubated with 0.01 µM Hb-488 in the presence or absence of 2 μM sSortilin-ALFA or 2 μM albumin for 16 h, and analyzed by FCM. (F). (G) Schematic of in vivo clearance of hemoglobin (Hb) after hemolysis. Mice were given 4 mg phenylhydrazine with 0.1 mg AF38469 or DMSO via intraperitoneal injection, and blood collected at 12, 24, 36, 48, and 72 h later. (H) Left panel: Plasma samples at the 72-h time point. Right panel: Effect of AF38469 on the PHZ-dependent increase in circulating Hb. The values are relative to the maximum level observed in the PHZ + DMSO group, which was set to 1. (I) Schematic of Hb in vivo clearance. Following intravenous injection of 60 µg Hb-biotin alone or with 170 µg sSortilin-Alfa, blood samples were collected at 1, 5, 10, 20, 30, and 60 min. (J) The Hb blood clearance kinetics in the presence of sSortilin. Relative changes in the plasma Hb-biotin level. The Hb-biotin concentration at 1 min post-injection was set to 100%. (K) A schematic model illustrating that sortilin facilitates the endocytosis of leaked cytosolic proteins. The relative net fluorescent intensity in FCM is the -fold change in the net fluorescent intensity of each sample normalized to the net fluorescent intensity of control cells (B, E), albumin-treated cells (C), or none cells (F), all of which were set to 1. Data are presented as means ± s.e.ms.; n = 3 (C, E, F, J), n = 4 (B), n = 9 (H). *P < 0.05.

The degradation of leaked cytosolic proteins is physiologically important, as illustrated by the need for hemoglobin (Hb) clearance. Hb, an intracellular, non-glycosylated protein, is released into the extracellular space during hemolysis, where its free iron can trigger toxic oxidative stress^49^. While the primary Hb clearance system involves haptoglobin and the CD163 receptor, both of which mediate lysosomal Hb degradation, haptoglobin- or CD163-KO mice exhibit only a slight reduction in leaked Hb degradation^50,51^, suggesting the involvement of an alternative clearance pathway. We therefore used Hb as a model non-glycosylated cytosolic protein (Figure 4D). We observed increased Hb internalization in sortilin-OE cells, which was inhibited by AF38469 and excess sSortilin (Figures 4E and F). To investigate whether Hb clearance is sortilin-dependent *in vivo*, we induced hemolysis in mice using phenylhydrazine (PHZ)^52^, with or without AF38469 (Figure 4G). Seventy-two hours after intraperitoneal PHZ injection, plasma from AF38469-treated mice exhibited a pronounced red color, indicating increased intravascular hemolysis (Figure 4H). Plasma Hb levels were higher in mice co-administered AF38469. To further characterize the extracellular function of sortilin, we co-injected biotinylated Hb (Hb-biotin) and sSortilin into the mouse tail vein (Figure 4I). The half-life of Hb-biotin was significantly prolonged when co-administered with sSortilin (Figure 4J). Together, these findings indicate that sortilin acts as a scavenger receptor for clearing leaked cytosolic proteins from the extracellular milieu (Figure 4K).

### A hydrophobic interaction is key to uptake

Unlike conventional receptors that recognize substrates through specific binding structures, sortilin is unlikely to form tailored interactions with diverse cytosolic proteins. Given that hydrophilic glycans often shield protein surfaces, we hypothesized that sortilin recognizes small exposed hydrophobic regions (Figure 5A). To investigate the role of hydrophobic interactions, we tested the effects of detergents. The binding of sSortilin to both RFP-GFP and deglycosylated ANGPT4 was significantly inhibited by the nonionic detergent dodecyl β-D-maltoside (DDM) and the zwitterionic detergent CHAPS (Figures 5B and C), indicating the crucial role of hydrophobic interactions in sortilin–substrate binding.

**Figure 5.**
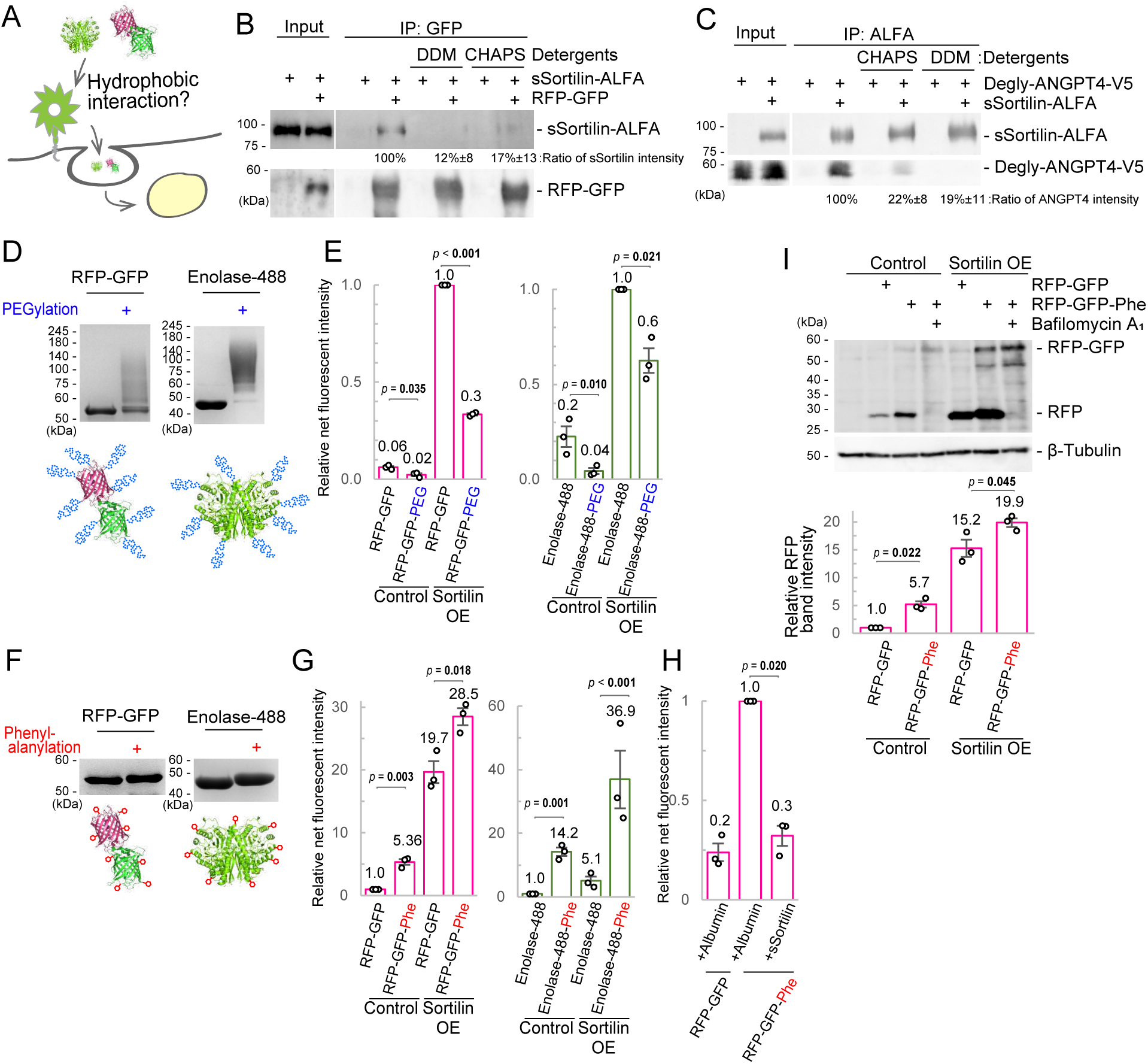
Non-glycosylated proteins bind to sortilin via hydrophobic interactions. (A) Schematic representation whether sortilin dependent endocytosis is mediated via hydrophobic interactions. (B and C) In vitro immunoprecipitation of sSortilin in the presence of detergents. sSortiln-ALFA-His, and RFP-GFP or deglycosylated ANGPT4-V5, were mixed and subjected to immunoprecipitation using anti-GFP or anti-ALFA nanobody beads in the presence or absence of 0.5% DDM or 0.5% CHAPS. The resulting precipitates were then immunoblotted using antibodies against GFP, V5, and ALFA. (D) PEGylation of RFP-GFP and enolase. Purified RFP-GFP and enolase were conjugated with 5-kDa NHS-PEG and subjected to SDS-PAGE followed by Coomassie Brilliant Blue staining. (E) FCM analysis of sortilin-dependent internalization of PEGylated proteins. Control and sortilin OE cells (HEK293) were incubated with 0.5 μM RFP-GFP, 0.5 μM RFP-GFP-PEG, 0.01 µM enolase-488, or 0.01 µM enolase-488-PEG, for 16 h, and analyzed by FCM. (F) Phenylalanylation of RFP-GFP and enolase. Purified RFP-GFP and enolase were conjugated with 5-kDa NHS-phenylananine and subjected to SDS-PAGE followed by Coomassie Brilliant Blue staining. (G) FCM analysis of sortilin-dependent internalization of phenylalanylated proteins. Control and sortilin OE cells (HEK293) were incubated with 0.5 μM RFP-GFP, 0.5 μM RFP-GFP-phenylalanine, 0.01 µM enolase-488, or 0.01 µM enolase-488-phenylalanine, for 16 h, and analyzed by FCM. (H) Competitive inhibition of phenylalanylated RFP-GFP internalization by sSortilin. HEK293 cells were incubated with 0.5 μM RFP-GFP-phenylalanine in the presence or absence of 2 μM sSortilin-ALFA or 2 μM albumin for 16 h, and analyzed by FCM. (I) Immunoblot analysis of RFP cleavage using RFP-GFP-phenylalanine. Control and sortilin OE cells (HEK293) were incubated with 0.5 μM RFP-GFP or 0.5 μM RFP-GFP-phenylalanine for 16 h. Relative RFP intensity is shown as the fold change relative to RFP-GFP-treated control cells, which was set as 1. The relative net fluorescent intensity in FCM is the -fold change in the net fluorescent intensity of each sample normalized to the net fluorescent intensity of sortilin OE cells (E), control cells (G), RFP-GFP-Phe with albumin treated cells (H), all of which were set to 1. Data are presented as means ± s.e.ms.; n = 3 (B, C, E, G, H, I).

Next, we mimicked glycosylation by conjugating polyethylene glycol (PEG), a hydrophilic polymer, to the primary amines of lysine residues on RFP-GFP and enolase-488 using NHS ester chemistry, which yielded approximately 6–10 molecules of 5-kDa PEG per protein molecule (Figure 5D). PEGylation of both RFP-GFP and enolase-488 significantly reduced internalization in control and sortilin-OE cells (Figure 5E). Furthermore, we performed the reciprocal experiment by covalently conjugating phenylalanine residues to the proteins via NHS ester chemistry to increase their surface hydrophobicity (Figure 5F). Consistent with our hypothesis, phenylalanine conjugation to RFP-GFP and enolase-488 markedly increased their sortilin-dependent endocytosis (Figures 5G and H). Immunoblotting of the RFP cleavage assay also showed that phenylalanine addition enhanced its lysosomal degradation (Figure 5I). Collectively, these results suggest that sortilin recognizes hydrophobic regions on the protein surface.

### Sortilin hydrophobic patches facilitate protein endocytosis

Reasoning that hydrophobic interactions facilitate substrate recognition, we hypothesized that sortilin contains surface HPs. To map these patches, we used AGGRESCAN3D to predict potential hydrophobic surface cores from the 3D structure of sortilin^53,54^. This analysis identified three HPs, which we named HP1–3. HP1 (L132–V135) and HP2 (F590–I591) are located on protruding loops, while HP3 (V498–V500) is situated on a blade surface (Figure 6A). Binding assays with phenyl sepharose, a hydrophobic matrix, revealed that sSortilin ΔHP123, a mutant lacking all three patches, had significantly lower binding affinity than sSortilin WT (Figure 6B), demonstrating that deletion of these patches decreases the surface hydrophobicity of sortilin. Furthermore, sSortilin ΔHP123 exhibited a significantly reduced binding affinity for deglycosylated ANGPT4-V5 (Figure 6C), suggesting that surface HPs on sortilin mediate its interaction with substrate proteins.

**Figure 6.**
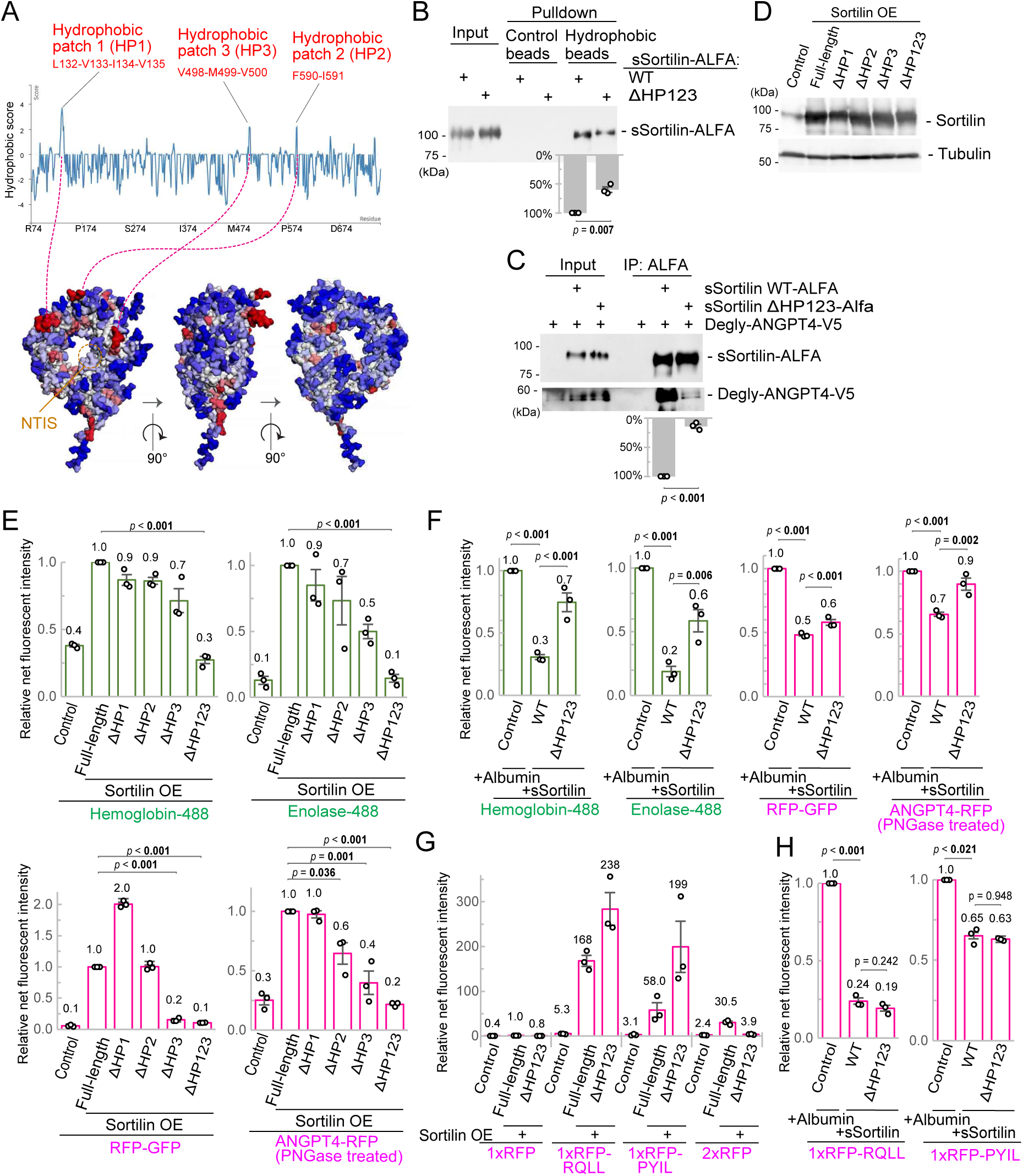
Hydrophobic patches on sortilin mediate protein endocytosis. (A) Three hydrophobic patches (HPs) identified using Aggrescan3D. (Top) Hydrophobicity score per residue of sortilin. (Bottom) 3D structure with hydrophobic (red) and hydrophilic (blue) amino acids. The green dotted circle indicates the neurotensin interaction site (NTIS). (B) Pulldown assay of sortilin with hydrophobic beads. sSortilin-ALFA-His WT or Δ HP123 was mixed with hydrophobic beads (Phenyl Sepharoses). The resulting precipitates were then subjected to immunoblotting using antibody against ALFA. (C) In vitro immunoprecipitation of sSortilin WT or ΔHP123 with ANGPT4. Deglycosylated ANGPT4-V5 and sSortilin-ALFA-His WT or ΔHP123 were mixed and subjected to immunoprecipitation using anti-ALFA nanobody beads. The resulting precipitates were then immunoblotted using antibodies against V5, and ALFA. The relative band intensity is the -fold change of each sample intensity, normalized to the intensity of the sample containing sSortilin WT, which was set to 100%. (D) Expression levels of sortilin full-length Δ HP1, ΔHP2, ΔHP3, and ΔHP123 mutant proteins. Lysates from the indicated cells were subjected to immunoblotting using antibodies against sortilin and β-tubulin. (E) FCM analysis of internalization by sortilin mutant OE cells. Control cells and the indicated sortilin OE cells (HEK293) were incubated with the listed proteins for 16 h, and analyzed by FCM. (F) Inhibition of sortilin-dependent internalization of various substrates by excess sSortilin. Control or sortilin mutant OE cells (HEK293) were incubated with the indicated proteins in the presence of 4 μM sSortilin-ALFA or albumin for 16 h, and analyzed by FCM. (G) NTIS-dependent uptake of sortilin ΔHP123 remains intact. HEK293 cells were incubated with the 0.5 μM of the listed proteins for 16 h, and analyzed by FCM. (H) sSortilin ΔHP123 inhibits NTIS-dependent uptake. HEK293 cells were incubated with 0.5 μM indicated proteins in the presence of 4 μM sSortilin-ALFA WT, sSortilin Δ HP123-ALFA, or albumin for 16 h, and analyzed by FCM. The relative net fluorescent intensity in FCM is the -fold change in the net fluorescent intensity of each sample normalized to the net fluorescent intensity of full-length sortilin OE cells (E), albumin-treated cells (F, H), and full-length sortilin OE cells in 1xRFP (G), all of which were set to 1. Data are presented as means ± s.e.ms.; n = 3 (B, C, D, E, F, G).

To determine the functional involvement of HPs in endocytosis, we established stable cell lines expressing sortilin ΔHP mutants at comparable levels (Figure 6D). While ΔHP1 and ΔHP2 had subtle, substrate-specific influences, ΔHP3 was critically required for efficient endocytosis, with ΔHP123 resulting in a dramatic loss of uptake capacity (Figure 6E). Consistent with this, the competitive inhibitory effect of sSortilin ΔHP123 was significantly reduced compared with that of sSortilin WT (Figure 6F). The C-terminal tetrapeptides of neurotensin and progranulin bind within the central cavity of sortilin—the neurotensin interaction site (NTIS)^55,56^—which is distinct from its HPs (Figure 6A). In agreement with previous studies showing that the C-terminal PYIL and RQLL motifs of neurotensin and progranulin, respectively, mediate sortilin-specific internalization^56,57^, both 1xRFP-PYIL and 1xRFP-RQLL exhibited greater cellular internalization than 1xRFP alone (Figure 6G). Notably, the ΔHP123 mutation did not decrease 1xRFP-PYIL and 1xRFP-RQLL uptake; rather, it enhanced it, demonstrating that ΔHP123 does not compromise overall receptor integrity. In addition, competitive inhibition of sSortilin ΔHP123 on the cellular uptake of 1xRFP-RQLL and 1xRFP-PYIL was equivalent to those of sSortilin WT (Figure 6H). Taken together, these results demonstrate that the HPs of sortilin constitute a novel binding domain, distinct from the canonical NTIS, that confers broad-spectrum recognition of cytosolic proteins for endocytic uptake.

### Rapid cycling of sortilin drives protein endocytosis

GFP-sortilin was scarcely detectable at the cell surface; instead, it was predominantly localized to endosomes and Golgi-like structures (Figure 7A), consistent with an earlier report^58^. In GFP-sortilin Tet-On cells, GFP-sortilin levels remained unchanged during 2xRFP substrate uptake (Figure S6A), suggesting that sortilin itself is not co-transported to lysosomes alongside its substrates. Together, these findings led us to hypothesize that sortilin operates via rapid cycling, stochastically shuttling extracellular substrates to the lysosome for degradation.

**Figure 7.**
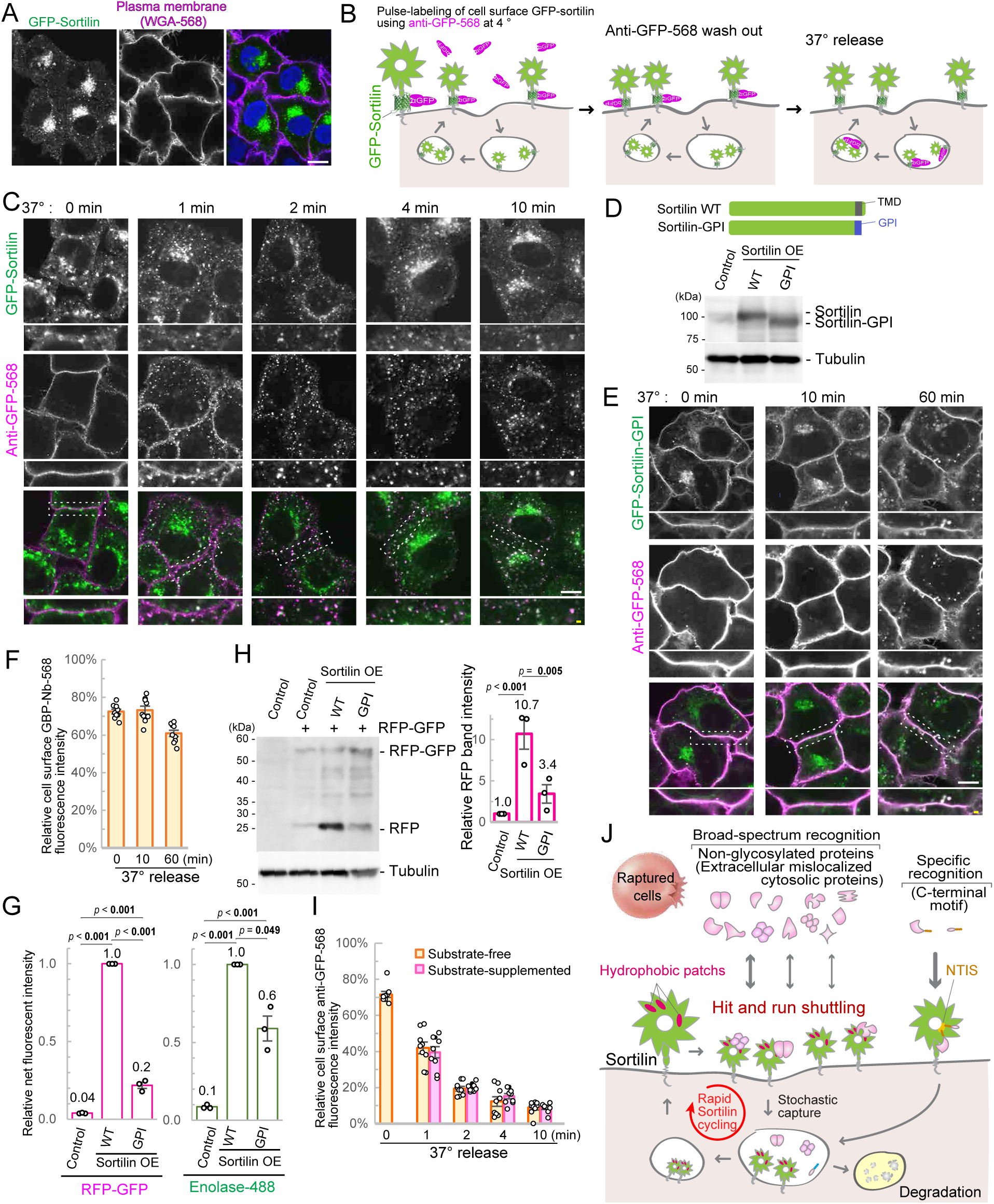
Sortilin facilitates endocytosis via hit-and-run shuttling. (A) Cellular localization of sortilin. HeLa cells stably expressing GFP-sortilin were fixed and stained with WGA-568 and observed by confocal microscopy. (B) Schematic representation of pulse-chase assay of cell-surface GFP-sortilin. Cell-surface GFP-sortilin was pulse-labeled with anti-GFP nanobody-ATTO-568 (anti-GFP-568) at 4°C for 30 min. After washing with PBS, cells were chased in medium at 37°C for the indicated time periods. (C) Pulse-chase assay of GFP-sortilin. HeLa cells stably expressing GFP-sortilin were selectively labeled at the cell surface with anti-GFP-568 at 4°C for 30 min, followed by a time-course chase at 37°C (0, 1, 2, 4, and 10 min) in serum-free DMEM (No serum) prior to confocal imaging. (D) Generation of sortilin-GPI-expressing cells. Lysates from the indicated cells (HEK293) were subjected to immunoblotting using antibodies against sortilin and β-tubulin. (E) Pulse-chase labeling and confocal imaging were performed as in (C), except that HeLa cells stably expressing GFP-sortilin-GPI were analyzed at 0, 10, and 60 min of chase in substrate-free condition. (F) Time-course quantification of surface GFP-sortilin-GPI internalization. Relative surface fluorescence intensity (cell periphery/total cell signal ratio of anti-GFP-568) is shown. Data represent mean ± SEM (n = 3 independent experiments, 9 fields of view in total). (G) FCM analysis of internalization by sortilin-GPI OE cells. Control cells and the indicated sortilin OE cells (HEK293) were incubated with the listed proteins for 16 h, and analyzed by FCM. The relative net fluorescent intensity in FCM is the -fold change in the net fluorescent intensity of each sample normalized to the net fluorescent intensity of control cells, all of which were set to 1. (H) Immunoblot analysis of the RFP-GFP cleavage assay data on control and sortilin-GPI OE cells (HEK293). Control and sortilin OE cells (HEK293) were incubated with 0.5 μM RFP-GFP for 16 h, and immunoblotted using anti-mCherry and -β-tubulin antibodies. (I) Time-course quantification of surface GFP-sortilin internalization with or without substrate. Cells were treated as in (F) in serum-free DMEM (substrate-free) or 10% FBS in DMEM (substrate-supplemented). Relative surface fluorescence intensity (cell periphery/total cell signal ratio) was calculated as described in (G). Data represent mean ± SEM (n = 3 independent experiments, 9 fields of view in total). (J) Proposed model for sortilin-mediated endocytosis via Hit-and-Run shuttling. Scale bars: 10 μm, 2 μm (inset).

To test this hypothesis, we conducted pulse-chase labeling of cell-surface GFP-sortilin using an ATTO-568-conjugated anti-GFP nanobody (anti-GFP-568) to trace its surface residence time and internalization kinetics (Figure 7B). Anti-GFP-568 specifically labeled the surface of GFP-sortilin–expressing cells, but not non-expressing cells (Figure S6B). GFP-sortilin was internalized within minutes during the chase under substrate-free conditions (serum-free DMEM) (Figure 7C), indicating that its cell surface residence time is less than 2 minutes.

To test whether increasing the surface residence time of sortilin compromises its efficiency in substrate endocytosis, we generated a sortilin mutant in which the transmembrane domain (TMD) was replaced with a GPI anchor signal (GFP-sortilin-GPI) (Figure 7D). Pulse-chase analysis revealed that GFP-sortilin-GPI exhibited a markedly increased surface residence time, with a substantial fraction remaining at the plasma membrane even after 60 min (Figures 7E and F). Quantitative assessments using FCM and the RFP cleavage assay showed that substrate internalization efficiency was significantly reduced in sortilin-GPI compared with sortilin WT (Figures 7G, 7H, and S6D). Notably, fluorescence microscopy revealed that substrates were internalized rather than merely accumulating at the cell surface, even in cells expressing sortilin-GPI (Figure S6C), indicating that modest endocytic activity was retained in cells expressing sortilin-GPI. These findings suggest that the cytoplasmic C-terminus is not required for substrate endocytosis; instead, sortilin physically co-internalizes with its substrates into endosomes through rapid sortilin cycling.

To determine whether sortilin internalization itself is substrate-independent, we monitored cell-surface GFP-sortilin levels in the presence or absence of substrates. Its residence time was completely unaffected by substrate presence (Figure 7I). Together, these findings indicate that sortilin does not rely on ligand-induced endocytosis but operates via constitutive cycling that efficiently clears diverse substrates through a “hit-and-run” shuttling mechanism (Figure 7J).

## Discussion

Intracellular PQC systems clear mislocalized organellar proteins within the cell^4–8^. We reveal a novel ePQC system, mediated by sortilin, which recognizes leaked cytosolic proteins that are mislocalized outside the cell, targeting them for lysosomal degradation. Specifically, our findings demonstrate that sortilin preferentially internalizes non-glycosylated substrates (Figure 3), drives uptake in a hydrophobicity-dependent manner (Figure 5), and recognizes substrates via HPs (Figure 6). The constitutive cycling of sortilin between the cell surface and endocytic compartments proceeds independently of substrate availability (Figure 7). The endocytosis of these cytosolic proteins is driven by a mechanism we term “hit-and-run” shuttling, which integrates the ability of sortilin to engage diverse substrates with its rapid receptor cycling (Figure 7J).

Compared with non-selective fluid-phase endocytosis, which randomly routes extracellular solutes to endosomes, sortilin operates as a sophisticated ePQC receptor that preferentially targets leaked cytosolic proteins for lysosomal degradation while largely sparing functional extracellular glycoproteins from clearance. Although the interaction between sortilin and certain substrates (e.g., RFP-GFP) is characterized by a rapid dissociation rate (t1/2 ∼1 min; Figure 1F), the kinetic balance with the residence time of sortilin at the cell surface (t1/2 ∼2 min; Figure 7I) allows a significant fraction (33%) of bound substrates to be internalized per binding event. Substrates with higher affinity are likely internalized more efficiently. Rather than fine-tuning specific affinities for each substrate, sortilin appears to function as a scavenger receptor through a simpler strategy: accommodating diverse substrates via HPs, coupled with rapid receptor cycling, thereby eliminating the need for tailored binding modes for individual substrates. Given the recent report that the sortilin substrate thyroglobulin preferentially interacts with monomeric rather than dimeric sortilin via the hydrophobic loop in blade 1 (corresponding to HP1 in this study)^59,60^, HP1 likely exerts a dual function. While mediating substrate binding at the cell surface, it may also drive sortilin dimerization in the acidic environment of endosomes, thereby acting as a structural safeguard to effectively suppress substrate re-binding.

We propose that the absence of glycosylation in the extracellular space acts as a danger signal marking abnormal proteins for clearance. This suggests a paradoxical relationship with intracellular glycosylation. In the cytosol, glycosylation serves as a danger signal: when an invading bacterium ruptures the endosomal membrane, the luminal-facing glycoproteins of that membrane become exposed to the cytoplasm and directly recognized by galectins. This triggers autophagy, which clears both the invading bacterium and the damaged endosome^13,61,62^. Similarly, glycosylated proteins retro-translocated from the ER to the cytosol are recognized by E3 ligases, ubiquitinated, and degraded by the proteasome^63,64^. By contrast, as most extracellular proteins are glycosylated, the lack of a glycan on a protein in the extracellular space can serve as a signature of abnormality, perhaps indicating a cytosolic origin, thereby promoting its elimination. Some secreted proteins, however, are naturally non-glycosylated, particularly smaller peptides such as hormones (e.g., insulin). We found that sortilin internalized 2xRFP with more than 10-fold greater efficiency than 1xRFP (Figure S1C). This may reflect increased avidity of protein binding, suggesting that smaller proteins may be less effectively recognized by sortilin. For example, while peptide hormones are not glycosylated, their small size may allow them to readily evade sortilin-dependent degradation. Although neurotensin and progranulin bind sortilin with high affinity by inserting their C-terminal four-residue motif into the tunnel of the ten-bladed β-propeller domain^55–57,65–67^, this NTIS does not overlap with the HPs defined here (Figure 6A). Indeed, 1xRFP-RQLL and 1xRFP-PYIL were markedly internalized into sortilin-OE cells in an HP-independent manner (Figure 6G). Taken together, our findings indicate that sortilin operates via a dual mode of substrate recognition: broad-spectrum recognition driven by surface HPs, and motif-specific recognition through the NTIS. AF38469 also binds to the NTIS of sortilin^37,55^, a site distinct from the outer HPs (Figure 6A). Meanwhile, the effective inhibition of sortilin-dependent substrate endocytosis by AF38469 (Figure 1H) is explained by direct steric hindrance^68^. Occupation of the inner tunnel by AF38469 may create a physical barrier that restricts the docking and proper spatial orientation of bulkier substrates. Alternatively, the NTIS might also contribute to the internalization of some proteins that lack specific recognition motifs.

Glycosylation of proteins extends their circulating half-life, a principle widely exploited to engineer protein therapeutics with enhanced stability, such as erythropoietin^69,70^. While protection from the hepatic asialoglycoprotein receptor (ASGPR) is conventionally cited to explain the extended half-life of glycosylated proteins^70^, the ASGPR model fundamentally concerns recognition of terminal glycans (sialic acid vs. exposed galactose) rather than the absolute necessity of glycosylation itself^71^. Indeed, ASGPR cannot account for the rapid clearance of completely non-glycosylated proteins. Our findings provide a long-sought mechanistic basis for this phenomenon: glycosylation physically shields hydrophobic protein surfaces from recognition by sortilin. This is supported by the finding that removal of all glycans by PNGase F was associated with a much greater increase in substrate uptake than other glycosidase treatments, including sialidase (neuraminidase A) (Figure S4D). Consistent with this, conjugation with hydrophilic PEG, which mimics the biophysical properties of glycans, similarly attenuated sortilin-dependent degradation (Figure 5E). Thus, glycosylation may at least partly extend protein half-life by allowing proteins to evade sortilin-mediated endocytic degradation, whereas sialic acid capping serves as a secondary timer against lectin-mediated clearance.

Importantly, our data also show that glycosylation does not confer complete protection against sortilin-facilitated turnover. Although a few glycans cannot fully shield the protein surface and so do not completely block degradation, they markedly reduce clearance efficiency (Figure 3B). Consistent with this, even glycosylated proteins exhibited increased degradation in sortilin-expressing cells (Figure 3). Together, these findings raise the possibility that sortilin is a major regulator of global extracellular protein turnover. In a functional parallel, the constant non-selective turnover of bulk cytosolic proteins via autophagy is believed to be vital for eliminating aberrant proteins that bypass selective degradation. Autophagic deficiency has been linked to the pathogenesis of neurodegenerative disease^14,72^. Notably, sortilin family members have been associated with neurodegenerative diseases, including Alzheimer’s disease^27^. Analogous to this cytosolic mechanism, the non-selective clearance of extracellular proteins by the sortilin family may be critical for preserving extracellular proteostasis. It remains unclear whether sortilin deficiency is associated with accumulation of aberrant proteins in the extracellular space. Sortilin KO mice do not exhibit phenotypes consistent with severe neurodegeneration^25^; in contrast, they have been reported to exhibit neuroprotective effects, particularly age-dependent rescue of sympathetic neurons attributable to reduced programmed cell death^73,74^. As our data suggest that SorLA also contributes to extracellular protein degradation (Figure 2), a more comprehensive understanding is needed. It is also possible that fluid-phase endocytosis and sortilin-mediated endocytosis act in a complementary manner to facilitate the bulk, non-selective degradation of extracellular proteins.

Renal clearance is also involved in extracellular protein elimination. While small blood proteins (<30 kDa) are filtered out by the glomerular filtration barrier (GFB), larger proteins may be more likely to harbor HPs, facilitating their removal via sortilin. Small proteins unrecognized by sortilin are instead eliminated by renal filtration, suggesting that sortilin-facilitated degradation and renal clearance function in a complementary manner to execute comprehensive ePQC.

Mechanistically, macroautophagy, chaperone-mediated autophagy, and microautophagy route internal cytosolic components to the lysosome^75^, whereas heterophagy disposes of externalized materials via endocytosis^76,77^. Because Sortilin targets cytosolic proteins that have escaped into the extracellular space, this selective degradation can be broadly viewed as an extension of autophagic self-clearance, even though its membrane trafficking mode is fundamentally heterophagic. Consequently, the Sortilin-mediated clearance of leaked cytosolic proteins can be conceptualized as a specialized form of ‘hetero-autophagy,’ operating at the interface of endocytosis and proteostasis.

The field of targeted protein degradation has recently expanded to include the selective degradation of extracellular proteins with tools such as LYTACs^78–80^. Notably, sortilin induced lysosomal degradation when a protein was tagged with only the four C-terminal amino acids of its ligand, progranulin^57^. These findings support our model that sortilin acts as a constitutive scavenger receptor for extracellular proteins and suggest that its binding properties could be exploited for targeted protein degradation.

## Acknowledgements

This work was supported by JSPS KAKENHI grants (nos. 20H03249, 22H04634, 23H04932, and 24K02018 to EI); the JSPS Program for Forming Japan’s Peak Research Universities (J-PEAKS) (grant no. JPJS00420230002 to E.I.); the JST FOREST Program (grant no. JPMJFR204N to E.I.); the Takeda Science Foundation (to E.I.); and the Research Support Program of the Institute for Advanced Academic Research (IAAR) at Chiba University (to E.I.). We thank Dr. Yoshitaka Tanaka (Kyushu University) for anti-LAMP1 antibodies and Dr. Takeharu Nagai (Osaka University) for pcDNA3 Gamillus. We are also grateful to Yae Sano and Hiroko Takano for their excellent technical assistance. We also acknowledge the use of Gemini (Google) and Claude (Anthropic) for language refinement, proofreading, during manuscript preparation. The authors reviewed all output and take full responsibility the content of this paper.

## Author contributions

S.S. performed the experiments on Sortilin- and SorLA-KO cells. K.K. performed the mouse experiments. S.B. performed the experiments on internalization of de-glycosylated proteins. M.K. performed preliminary evaluation of the endocytosis assay using fluorescent proteins. A.M. provided critical advice. E.I. conceived the study, designed the experiments, performed most of the experiments and analysis, wrote the original draft, acquired funding for the study, and supervised the work. All authors contributed to paper review and editing.

## Methods

### Cell culture

Flp-In T-Rex HEK293 [Thermo Fisher Scientific (TFS), cat#R78007], HepG2 (RIKEN BRC, cat#RCB1648), HeLa (RIKEN BRC, RCB0191), and U2OS (kindly provided by Dr. Taka-aki Tamura, Chiba university, Japan) cells were cultured in Dulbecco’s Modified Eagle’s Medium (DMEM, Nacalai Tesque). T98G (Japanese Collection of Research Bioresources (JCRB, cat#IFO50303) and U251-MG (JCRB, cat#IFO05288) cell lines were maintained in Eagle’s Minimum Essential Medium (MEM, Nacalai Tesque). HuEhT-2 cells (JCRB, cat#JCRB1459) were cultured in MCDB131 medium (TFS) with 0.03 g/L endothelial cell growth supplement (ECGS), 5 µg/mL heparin, and 10 mM L-glutamine. RAW 264 (RIKEN BRC, cat#RCB0535) cell lines were maintained in Minimum Essential Medium (MEM, Nacalai Tesque) with 0.1 mM non essential amino acids (NEAA). J774.1 cell linnes (JCRB, cat#JCRB0018) were mintained in Roswell Park Memorial Institute 1640 medium (RPMI1640, Nacalai Tesque). All culture media were supplemented with 10% fetal bovine serum (FBS, Biosera) and 50 µg/mL penicillin-streptomycin. Cell growth proceeded in a humidified incubator at 37°C under 5% carbon dioxide. To generate stable cell lines expressing doxycycline (Dox)-inducible secreted proteins (e.g., sortilin), Flp-In T-Rex HEK293 cells were co-transfected with plasmids encoding the respective proteins and pOG44, which encodes the FLP recombinase. Transfected cells were selected and maintained using 100 µg/mL hygromycin. Gene expression at the FRT site was induced with 200 ng/mL Dox (Clontech).

### Plasmids

To construct pcDNA5 FRT TO sortilin (mouse full-length Sortilin), pcDNA5 FRT TO sSortilin-Alfa [SortilinΔTMD(1-753aa)-Alfa-6xHis] and pcDNA5 FRT TO sortilin-sfGFP (sfGFP inserted before Val755) mouse Sortilin cDNAs were PCR-amplified from total mouse Min6 cell cDNA. To construct pcDNA5 FRT TO SorCS1, SorCS2, SorCS3, or SorLA, cDNAs were amplified from the cDNA clone vectors (DNAFORM 40114197 for SorCS1, DNAFORM 100063568 for SorCS2, Flexi ORF clone ORK00727 for SorCS3, Flexi ORF clone ORK02134 for sorLA). To construct pcDNA5 FRT TO ANGPT4-mCherry(−6xHis), ANGPT4-V5(−6xHis), human ANGPT4 DNA was synthesized using gBlock Gene Fragments (Integrated DNA Technologies; Coralville, IA) and the amplified cDNA inserted into the pcDNA5 FRT TO vector along with mCherry, sfGFP, ALFA-tag, V5-tag and 6xHis-tag sequences. Sortilin deletion mutants [mouse Sortilin ΔHP1(deletion 132-135 aa), ΔHP2 (deletion 590-591 aa), ΔHP3 (deletion 498-500aa) and ΔHP1+2+3 (ΔHP123)] were generated via inverse PCR and Gibson assembly using the pcDNA5 FRT TO sortilin vector. Sortilin-GPI was constructed by fusing amino acids 1–753 of sortilin with the GPI anchor signal sequence (amino acids 228–253) of human PrP. pOG44 was used for the Flp-In system. To construct pLenti-puro Sortilin, mouse sortilin was inserted into pLenti CMV GFP Puro (Addgene 17448). To construct pRSETA RFP-GFP (mCherry-sfGFP-6xHis), pRSETA 2xRFP (mCherry-mScarleti-6xHis), pRSETA 1xRFP (mCherry-6xHis), 1xRFP (6xHis-mCherry),pRSETA 1xRFP-RQLL (6xHis-mCherry-RQLL), pRSETA 1xRFP-PYIL (6xHis-mCherry-PYIL), pRSETA 2xGamillus (Gamillus-Gamillus-6xHis), and pRSETA 1xGamillus (Gamillus-6xHis), the required DNAs were amplified from pCW57.1 The lysosomal-METRIQ vector (Addgene: 135401) containing sfGFP and mCherry, pmScarleti-C1(Addgene 85044), and pcDNA3 Gamillus (a gift from Dr. Takeharu Nagai, Osaka University)^34^. The DNA fragments were then inserted into the pRSET-A vector along with the ALFA-tag and 6xHis-tag sequences. pCMV-VSVG (Addgene, Cat#8454) and psPAX2 (Addgene, Cat#12260) were used for lentivirus production.

### Lentivirus infection

HEK293FT cells were transiently co-transfected with the lentiviral expression vector pLenti-puro-Sortilin, psPAX2, and pCMV-VSV-G using PEI MAX reagent (Polysciences). Four hours after transfection, the culture medium was replaced with fresh medium. After 72 h of incubation, the viral supernatant was harvested. HeLa cells were transduced with the collected viral supernatant for 24 h. Uninfected cells were subsequently eliminated by incubation with 1 µg/mL puromycin (InvivoGen).

### Generation of KO cells

KO cells were generated as decribed previously^18^. For sortilin and SorLA KO cells, human sortilin sgRNA (5ʹ-CATGTGTTTGATGATCTCAG-3ʹ) or human SorLA sgRNA (5ʹ-ACGCTTATGCCCAGTACCTC-3ʹ) was cloned into the lentiCRISPRv2 hygro vector (Addgene, cat#98291). As a control cells, control sgRNA (gControl)(5ʹ-CGCAGTCATTCGATAGGAAT-3ʹ) was used. Flp-In Trex-293 cells were infected with lentivirus expressing Cas9 and guide RNA. After 24 h, the cells were cultured in media with 100 µg/mL hygromycin for > 7 days. Single-cell clones were isolated via limiting dilution and KO cell lines subsequently confirmed by immunoblotting.

### Antibodies

Rabbit polyclonal anti-LAMP1 antibody was kindly provided by Y. Tanaka (Kyushu University, Fukuoka, Japan). Rabbit polyclonal anti-TRAPα antibody were a gift from R.S. Hegde (MRC LMB, UK). Rabbit polyclonal anti-GFP antibody, polyclonal anti-mCherry antibody, and polyclonal anti-ALFA tag antibody were generated by Eurofins using the full-length mCherry protein and the ALFA peptide as antigens in rabbits. Rabbit polyclonal anti-β-actin antibody (cat#20536-1-AP), rabbit polyclonal anti-sortilin antibody (cat#12369-1-AP) (for detection of sSortilin), rabbit polyclonal anti-SorCS1 antibody(cat# 23002-1-AP), and rabbit polyclonal anti-PEX14 antibody (cat# 10594-1-AP) were purchased from Proteintech. Rabbit polyclonal anti-sortilin antibody (cat# 20681S 12369-1-AP) (for detection of full-length sortilin) and the HRP-conjugated anti-rabbit antibody (cat#7074S) were purchased from Cell Signaling Technology (CST). Mouse anti-V5 antibody(cat# M215-3) was purchased from MBL. Rabbit polyclonal anti-SorLA antibody (cat# ab190684) was purchased from Abcam. Rabbit polyclonal anti-SorCS2 antibody (cat# AF4238) and anti-SorCS3 antibody (cat# MAB3067) were purchased from R&D. Mouse monoclonal anti-HSP90 antibody (cat# 610419) was purchased from BD biosciences. Rabbit polyclonal anti-Tom20 antibody (cat# sc-11415) was purchased from Santa cruz biotechnology.

ALFA nanobody beads were prepared by conjugating ALFA nanobody protein^81^ purified from pRSET-A ALFA-nanobody-6xHis to N-hydroxysuccinimide (NHS)-activated Sepharose 4 Fast Flow (GE). Similarly, GFP nanobody beads were prepared by conjugating GFP nanobody protein encoded by the the pOPINE GFP nanobody plasmid (Addgene #49712) to NHS-activated Sepharose 4 Fast Flow.

### Recombinant proteins from *Escherichia coli*

RFP-GFP (mCherry-sfGFP-6xHis), 2xRFP (mCherry-mScarleti-6xHis), and 2xGamillus (Gamillus-Gamillus-6xHis) received the 6xHis-tag and were then cloned into pRSET-A. Plasmids were introduced into the BL21(DE3) LOBSTR strain of *Escherichia coli* ^82^, and expression then induced with 0.3 mM isopropyl β-D-1-thiogalactopyranoside at 16°C. Cells were suspended in PBS with 300 mM NaCl and 10 mM imidazole, and disrupted by sonication. After ultracentrifugation, recombinant proteins in soluble fractions were purified using HisPur Cobalt Resin (TFS, cat# 89965). Purity was assessed via SDS-PAGE followed by Coomassie Brilliant Blue staining.

### Recombinant proteins secreted from mammalian cells

To encode proteins secreted from mammalian cells, pcDNA5 FRT TO plasmids containing sSortilin-ALFA [SortilinΔTMD(1-753aa)-Alfa-6xHis], ANGPT4-V5(−6xHis), or ANGPT4-mCherry(−6xHis) were introduced into the FLP site of the Flp-In T-Rex293 cell genome to generate stable cell lines (Tet-On). Secreted proteins in conditioned medium were purified as previouly described^83^. Briefly, cells were cultured for 4 days in serum-free advanced DMEM/F12 medium (TFS, cat#12634010) supplemented with L-glutamine and 0.2 µg/mL doxycycline (Dox). Each culture supernatant was centrifuged at 2,360 × *g* for 10 min at 4°C to remove dead cells. Secreted proteins were purified from the supernantant via Ni-NTA affinity chromatography and stored in PBS with 10% glycerol at 4°C.

### Fluorescent labeling of proteins

To generate albumin-565, ALT-488, enolase-488, GFP-Nanobody-ATTO-568 (anti-GFP-568), Hb-488, IgE-488, and cytosolic proteins-488, albumin (Fujifilm, Cat# 017-15146), alanine aminotransferase (ALT) (Fujifilm, Cat# 305-51931), enolase (Fujifilm, Cat# 301-50073), GFP nanobody protein, γ-glutamyltransferase (γGT) (Fujifilm, Cat# 305-50674), Hb (Sigma, Cat# H7379), IgE (Abcam, Cat# ab65866), and mouse tissue-derived cytoplasmic proteins (homogenized soluble lysates of liver, muscle, and heart, as well as red blood cells) were fluorescently labeled. One hundred micrograms of protein (or 1 mg of mouse tissue lysate) were/was mixed with 0.125 mg ATTO 488-NHS-Ester or 0.125 mg ATTO 565-NHS-Ester (ATTO-TEC) and incubated at 24°C for 1 h. The reaction was terminated by separation of the labeled protein from unreacted dye via Sephadex G-25 chromatography in PBS. Amyloid-β (1-42)-488 was purchased from Anaspec (cat# AS-60479-01). Transferrin-488 was purchased from Fujifilm (cat# 009-540-050). IgG-488 was purchased from TFS (cat# A-11001).

### PEGylation and phenylalanylation of proteins

To generate PEGlysated or phenylalanylated proteins, 1 mg of RFP-GFP or enolase-488 were mixed with 2.5 mg MeO-PEG-NHS (Iris Biotech, cat# PEG1165.0001) or N-t-Butoxycarbonyl-L-phenylalanine N-Hydroxysuccinimide Ester (Fujifilm, cat# 352-12741) and incubated at 24°C for 1 h. The reaction was terminated via separation of the labeled protein from unreacted compounds by Sephadex G-25 chromatography in PBS.

### Deglycosidase treatment

Fifteen units of PNGase F (NEB, cat# P0704S), 0.24 U β1-4 Galactosidase S (NEB, cat# P0745S), 0.15 U β-N-Acetylhexosaminidasef (NEB, cat# P0721S), or 0.6 U α2-3,6,8,9 Neuraminidase A (NEB, cat# P0722S) were mixed in 5 μL PBS with recombinant proteins (final concentration 10 μM ANGPT4-RFP, 10 μM ANGPT4-V5, 100 ng/μL IgG-488, 15 ng/μL IgE-488), and incubated at 37°C for 3 h. The recombinant proteins thus obtained were used in i endocytosis assays or immunoprecipitation.

### Endocytosis assay

Recombinant RFP-GFP (final concentration 0.2–0.5 μM), 2xRFP (final concentration 0.2–0.5 µM), 2xGamillus (final concentration 0.2–0.5 µM), ANGPT4-RFP (final concentration 0.2 µM), pHrodo-Green Dextran 10-kMW (final concentration 0.2 ng/ul), enolase-488 (final concentration 0.01 µM), ALT-488 (final concentration 0.01 µM), rGT-488 (final concentration 0.01 µM), albumin-488 (final concentration 0.03 μM), albumin-565 (final concentration 0.05–0.1 μM), amyloid-β-488 (final concentration 0.1–0.2 μM), IgG-488 (final concentration 2 ng/μL), IgE-488 (final concentration 0.3 ng/μL), hemoglobin-488 (final concentration 0.01 µM), or transferrin-488 (final concentration 5-10 ng/μL) were added to 250 μL of serum-free advanced DMEM/F12 (TFS, cat# 12634010)), a formulation that lacks serum but contains essential proteinaceous factors—such as 400 μg/mL (6 μM) albumin, 10 μg/mL (1.7 μM) insulin, and 7.5 μg/mL (0.1 μM) transferrin—to prevent nutrient starvation. Cells (25–50% confluency) in 24-well plates were washed with serum-free DMEM and then cultured with medium containing recombinant proteins at 37°C for 16 h. To assay competitive inhibition, sSortilin-ALFA (final concentration 1–4 μM), albumin (final concentration 0.5–500 μM), or AF38469 (Cayman, cat# 35530) (final concentration 0.3–10 µM) was mixed into the medium. In indicated experiments, 0.2 µM bafilomycin A_1_ (LC Laboratories, cat#B-1080) was added to the medium. The cells were subjected to flow cytometry, immunoblotting, or immunostaining.

### Flow cytometry (FCM)

Cells were detached with trypsin and resuspended in a solution containing 5% newborn calf serum and 1 µg/mL 4′, 6-diamidino-2-phenylindole (DAPI) in PBS. The cell suspension was then passed through a 70-µm cell strainer into a 1.5-mL tube and the collected cells subjected to flow cytometry using a CytoFLEX S platform (Beckman Coulter) equipped with 375-nm (for DAPI using the DAPI filter), 488-nm (for GFP and ATTO-488 using the FITC filter), and 561-nm (for mCherry and ATTO-565 using the ECD filter) lasers. Dead cells were detected via DAPI staining and excluded from analysis. Gating was sequential. First, cell populations were isolated based on the FSC-A and SSC-A scatterplots, followed by exclusion of cell aggregates using SSC-W vs. SSC-A or SSC-H to select for singlet cells. Dead cells were next eliminated by removing DAPI-positive cells. Finally, the green or red fluorescence intensity was measured in the remaining live singlet cells (total 10,000 cells). The net fluorescence intensity was determined by subtracting the mean fluorescence intensity (MFI) of the non-treated cells (auto-fluorescence) from that of cells of the endocytosis assays in the respective FITC or ECD channels. The relative net fluorescent intensity is the -fold change in the net fluorescent intensity of each sample, normalized to the net fluorescent intensity of control cells (indicated in the Figures), which was set to 1. Similarly, the relative fluorescence intensity was calculated by normalizing the MFI to that of the control cells (indicated in the Figures), which was set to 1.

### Cell lysis and immunoblotting

Cells were washed with cold PBS and lysed for 15 min at 4°C in lysis buffer (1% Triton-X100, 50 mM Tris/HCl, pH 7.5, 1 mM EDTA, 150 mM NaCl) containing a protease inhibitor cocktail (EDTA-free: Nakalai Tesque, cat#03969-21) and 1 mM phenylmethanesulfonyl fluoride (PMSF). After centrifugation at 20,000 × *g* for 5 minutes, each supernatant was collected, mixed with 6x SDS sample buffer, and boiled for 5 min at 95°C. Proteins were separated by SDS-PAGE, transferred to a polyvinylidene difluoride (PVDF) membrane (Millipore, cat#IPVH00010), and probed with antibody solutions diluted in Signal Enhancer HIKARI (Nakalai Tesque, cat#02270-81). Protein bands were visualized using ImmunoStar Zeta (Fujifilm Wako Pure Chemical Industries, cat#297-72404).

### Immunoprecipitation (IP)

For IP of RFP-GFP and sSortilin-ALFA, 125 nM RFP-GFP and 500 nM sSortilin-ALFA (for GFP IP), or 500 nM RFP-GFP and 125 nM sSortilin-ALFA (for ALFA IP), in 20 µL PBS, were pre-treated with 0.1 mM DSP for 30 min at 37°C and each sample then mixed with 230 µL DSP stop buffer (50 mM Tris-HCl 7.5, 0.01% BSA in PBS).

For IP of ANGPT4, PNGase-treated or -untreated ANGPT4-V5 was used. Purified 50 nM sSortilin-ALFA and 200 nM ANGPT4-V5 were mixed in 50 µL PBS with or without 0.5% CHAPS or 0.5% DDM. GFP nanobody or ALFA nanobody beads (5-µL bed volume) were added to the mixture followed by rotation at 4°C for 1 h. The beads were washed four times with PBS without 0.5% CHAPS or 0.5% DDM, transferred to new tubes, mixed with SDS sample buffer, and boiled for 5 min.

For pulldown assays using hydrophobic beads, 40 nM sSortilin-ALFA (WT or ΔHP123) in 50 µL of PBS was mixed with Phenyl Sepharose 6 Fast Flow (GE Healthcare, Cat# 17097310; 5-µL bed volume) and rotated at 4°C for 1 h. The beads were washed four times with PBS, transferred to new tubes, mixed with SDS sample buffer, and boiled for 5 min.

### Immunostaining

Cells cultured on cover slips were fixed at room temperature for 15 min with 3.7% formaldehyde in PBS. Prior to immunostaining, fixed cells were permeabilized for 5 min with 50 µg/mL digitonin in PBS. After blocking with 10% newborn calf serum in PBS for 30 min, cells were incubated for 1 h with the primary antibody. After washing three times with PBS, cells were incubated for 1 h with Alexa Fluor 568-conjugated goat anti-rabbit IgG antibody (TFS, cat# A-11011). For plasma membrane staining, cells were incubated with 1 µg/mL CF568-cojugated wheat geam agglutinin(WGA) (Biotium, Cat# 29077-1) for 30 min. Cells were then mounted with DAPI/DABCO/Glycerol/PBS. Stained cells were observed under a confocal laser microscope (FV1000IX81, Olympus) with a 100-x oil immersion lens (NA 1.40).

### Surface GFP-Sortilin Pulse-Chase Assay

HeLa cells stably expressing GFP-sortilin or GFP-sortilin-GPI were washed with cold Hank’s Balanced Salt Solution (HBSS) and pre-chilled at 4°C for 5 min. To label cell-surface GFP-Sortilin, cells were incubated with 5 µM anti-GFP nanobody conjugated to ATTO-568 (GFP-Nb-568) in HBSS at 4°C for 30 min. After washing with PBS to remove unbound nanobodies, internalization was initiated by incubating the cells at 37°C in either serum-free DMEM or DMEM supplemented with 10% FBS. At specified time points (0, 1, 2, 4, 10, and 60 min), the internalization was halted by washing with ice-cold PBS. Cells were fixed with 3.7% formaldehyde for 15 min, washed with PBS, and imaged using a confocal laser scanning microscope.

### Ethical Approval

All animal experimental procedures were performed in accordance with specified guidelines for the care and use of laboratory animals and were approved by the Animal Care and Use Committee of Chiba University(7-179).

### In vivo clearance experiments

In the hemolysis experiment, phenylhydrazine (PHZ) (Sigma, cat#114715) (4 mg per 20 g body weight) with DMSO or 0.1 mg of AF38469 were intraperitoneally injected into BALB/c mice (8 weeks of age). AF38469 was re-administered 36 hours after initial administration.

Recombinant Hb was labeled with NHS-biotin (Vector Laboratories cat# QUA10205) to generate Hb-biotin. BALB/c mice (8 weeks of age) were injected intravenously with 60 µg Hb-biotin with or without 170 µg sSortilin-Alfa protein per 20 g body weight.

Serial blood samples were taken via the tail vein at timed intervals. Four-microliter amounts of blood were mixed with 10 µL buffer (0.05 M EDTA in PBS) and plasma obtained from the anticoagulated blood via microcentrifugation at 1,200 x *g* for 15 min. The plasma was mixed with 2x SDS sample buffer and boiled for 5 min at 95°C. One microliter of each plasma sample was subjected to SDS-PAGE and proteins were detected using an anti-biotin HRP-linked antibody (CST, cat# 7075S) and an anti-Hb-beta antibody (Proteintech, cat# 16216-1-AP).

### Surface plasmon resonance (SPR)

SPR experiments were performed using a Biacore 1K system (Cytiva) at 25°C. Recombinant sSortilin was conjugated with Biacore cap-tag using the Biacore Cap-Tag Capture Kit (Cytiva) according to the manufacturer’s instructions and reversibly captured onto a Series S Sensor Chip CAP (Cytiva). HBS-EP buffer was used as the running and dilution buffer. To determine the binding kinetics, single-cycle kinetics analysis was conducted. Increasing concentrations of RFP-GFP (2.5, 5, 10, 20, and 40 µM) or enolase (0.05, 0.15, 0.5, 1.5, and 5 µM) (analyte) were sequentially injected over the sSortilin-captured surface without regeneration between injections, using default instrument parameters. A blank flow cell without sSortilin capture served as a reference surface. Sensorgrams were double-referenced by subtracting responses from the reference flow cell and blank buffer injections. Kinetic rate constants (k_on_ and k_off_) and the equilibrium dissociation constant (K_D_) were calculated by fitting the double-referenced sensorgrams to a 1:1 binding model using Biacore Insight Evaluation Software.

### Subcellular Fractionation

Mouse tissues (liver, skeletal muscle, and RBC) were pulverized in liquid nitrogen using a mortar and pestle. The frozen tissue powder was transferred to a homogenizer and suspended in five volumes of homogenization buffer consisting of PBS supplemented with 1 mM PMSF and 1× protease inhibitor cocktail. The tissue suspension was thoroughly homogenized on ice. The homogenate was centrifuged at 1,100 × *g* for 10 min at 4°C to remove nuclei and cellular debris, and the resulting supernatant was collected as the post-nuclear supernatant (PNS). The PNS was further centrifuged at 500,000 × *g* for 30 min at 4°C to separate the cytoplasmic and membrane fractions. The resulting supernatant was collected as the cytoplasmic fraction, aliquoted, and stored at −80°C. The remaining pellet was washed once with PBS and centrifuged again at 100,000 rpm for 30 min at 4°C. The washed pellet was resuspended in SDS sample buffer.

### Statistical Analysis

Data were collected from at least three biological replicates (N ≥ 3). Background-subtracted values were used for statistical testing in FCM (relative net fluorescent intenstiy) and immunoblot analyses. For relative fluorescent intenstiy of FCM, fluorescent values were used for statistiacal analysis. Prior to statistical testing, data were log-transformed with the exception of non-log-transformed data used in Figure 1D, 4H, and 4J. Statistical significance for multiple group comparisons was determined using repeated-measures (RM) ANOVA with Dunnett’s post hoc test (Figures 1C, 1D, 1H, 2C, 2E, 2G, 3B (for IgE-488), 3G, 4E, 4F, 5H, 6E, 6F, and 7H; and S1A, S2A, S2B, S2D, S2E, S3C, and S4A) or Tukey’s post hoc test (Figures 2F, 3H, and 7G; S1B, S1C, S2C, 3D, 4D, and 6D). Two-sample comparisons were analyzed using a paired two-tailed *t*-test (Figures 1E, 3B [for IgG-488], 3C, 3F, 4B, 4C, 4H, 4J, 5E, 5G, 5I, 6B, and 6C; and S1C), or Welch’s *t*-test (Figures 4H and 4J). Data in graphs are presented as relative values normalized to the control cells (set to 1.0), as indicated in the respective figure legends.

