## Supplemental figures for "Sortilin is a scavenger receptor for cytosolic proteins mislocalized to the extracellular milieu"

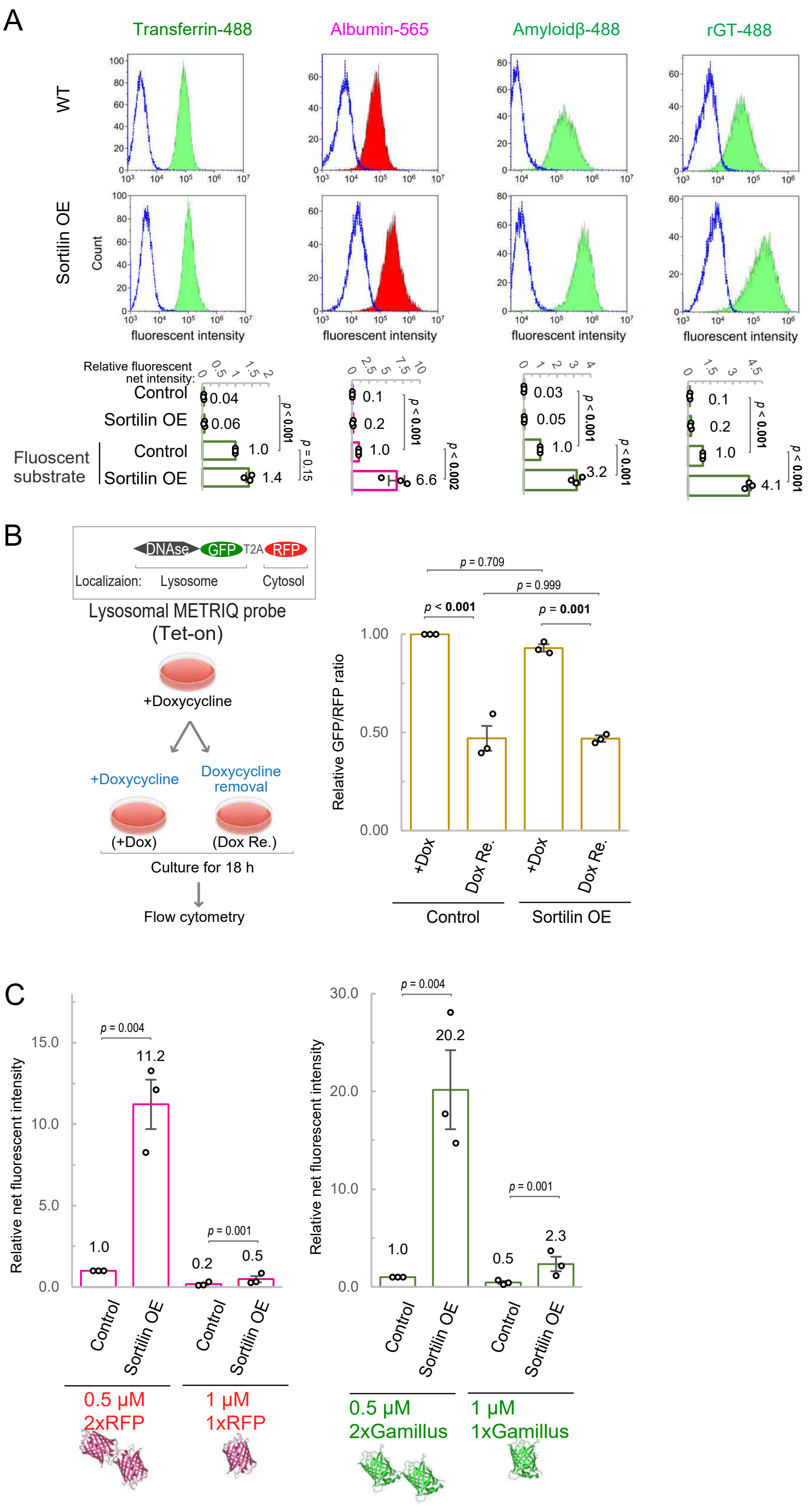


**Figure S1. Protein substrate specificity of sortilin-dependent internalization.**
(A) FCM analysis of substrate endocytosis as in Figure 1C with various substrates. Control and sortilin OE cells (HEK293) were incubated with the indicated proteins for 16 h, and analyzed by FCM. Sortilin-dependent transferrin internalization was significantly lower than that of other proteins (Figure. 1C). The blue dashed line in FCM data represents the signal from untreated control cells, and thus the level of cellular autofluorescence. (B) Lysosomal METRIQ analysis of sortilin OE cells. Left panel: A schematic of lysosomal METRIQ assay. Lysosomal METRIQ (Tet-On) cells (HEK293) were initially treated with doxycycline for several days to induce expression. Then, the cells were incubated for an additional 18 h with or without Dox. While GFP degradation proceeds normally when lysosomes are functional, its clearance is significantly prolonged upon the inhibition of lysosomal activity. Right panel: FCM analysis of lysosomal-METRIQ. The reduction in GFP levels remained fully intact in sortilin OE cells. (C) FCM analysis of size-dependent internalization. Control and sortilin OE cells (HEK293) were incubated with the indicated fluorescent proteins for 16 h, and analyzed by FCM. Although the number of fluorescent protein molecules was the same, 2xFP internalization was enhanced over 10-fold compared to that of 1xFP. In the 2xFP group, the control and Sortilin OE cells were identical to those of Figure 1C. The relative (net) fluorescent intensity in FCM is the -fold change of the (net) fluorescent intensity of each sample normalized to the (net) fluorescent intensity of control cells, all of which were set to 1. Data are presented as means ± s.e.ms.; n = 3 (A-C).

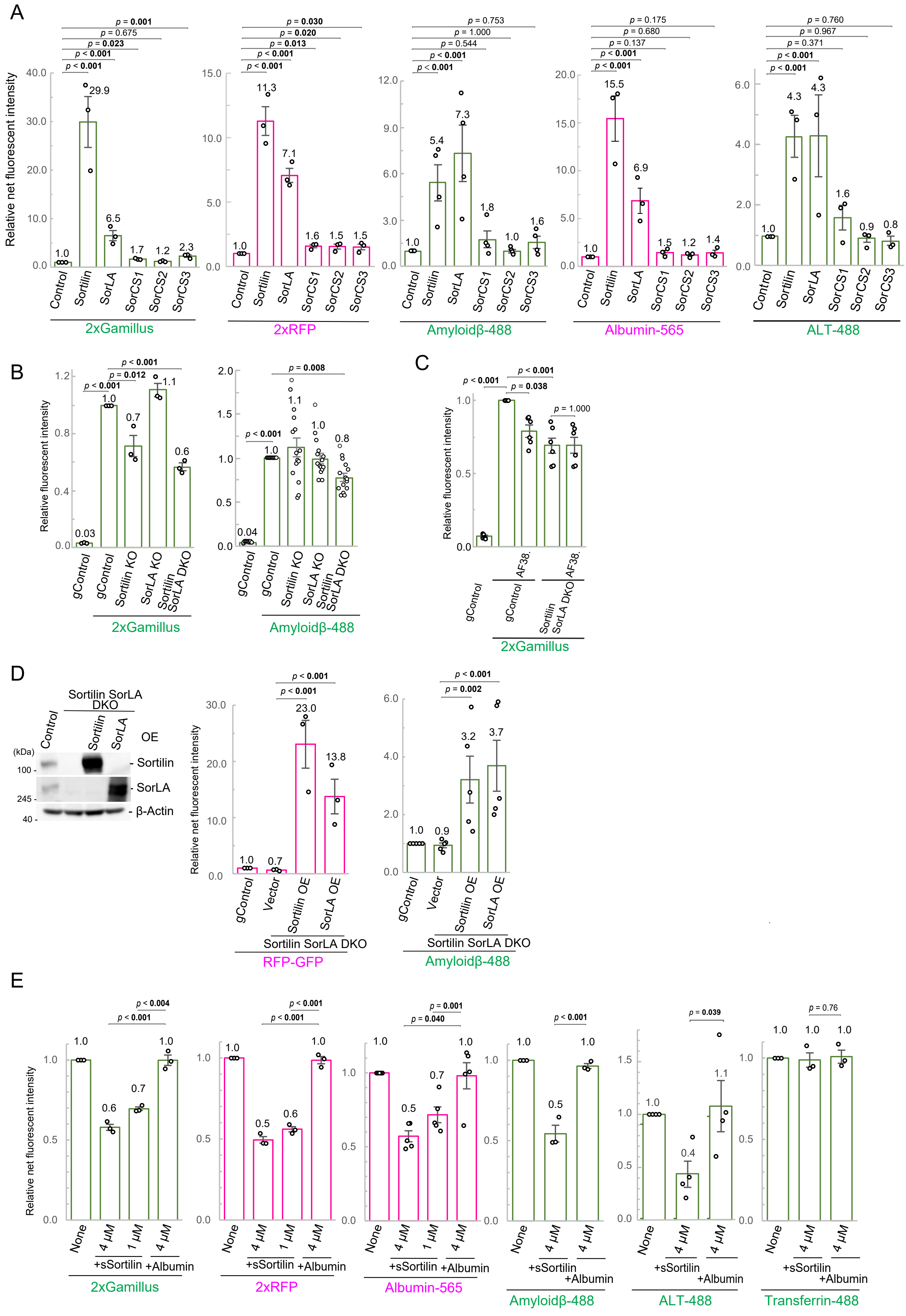
A
Relative net fluorescent intensity
Control
Sortilin
SorLA
SorCS1
SorCS2
SorCS3
2xGamillus
2xRFP
Amyloidβ-488
Albumin-565
ALT-488
B
Relative fluorescent intensity
gControl
Sortilin KO
SorLA KO
Sortilin SorLA DKO
C
AF38.
D
Sortilin SorLA DKO
OE
(kDa)
Sortilin
SorLA
β-Actin
Vector
Sortilin OE
SorLA OE
RFP-GFP
E
None
4 μM
1 μM
+sSortilin
+Albumin
Transferrin-488

Figure S2

**Figure S2. Figure S2. Sortilin and SorLA are involved in internalization of various extracellular proteins.**
(A) FCM analysis of fluorescent extracellular protein internalization in control and sortilin family OE cells. Control and indicated OE cells (HEK293) were incubated with the indicated proteins for 16 h. Fluorescent intensities per cell were measured by FCM. (B) FCM analysis of fluorescent extracellular protein internalization by control and KO cells. The control and indicated KO cells (HEK293) were incubated with indicated proteins for 16 h. Fluorescent intensities per cell were measured by FCM. (C) gControl or sortilin and SorLA DKO (HEK293) cells incubated with 0.5 μM 2xGamillus in the presence or absence of 10 μM AF38469 for 16 h and analyzed by FCM.
(D) Rescue of sortilin and SorLA DKO cells. Left panel: Sortilin or SorLA was overexpressed in sortilin and SorLA DKO cells. Right panel: FCM analysis of fluorescent extracellular protein internalization in DKO and rescued cells. gControl, DKO, and the indicated rescued cells (HEK293) were incubated with indicated proteins for 16 h. Fluorescent intensities per cell were measured by FCM. (E) Competitive inhibition of fluorescent protein internalization by sSortilin. HEK293 cells were incubated with indicated proteins in the presence or absence of 1 or 4 μM sSortilin-ALFA or 4 μM albumin for 16 h. Fluorescent intensities per cell were measured by FCM. The relative (net) fluorescent intensity is the -fold change of the (net) fluorescent intensity of each sample normalized to the net fluorescent intensity of control cells (A), gControl cells (B, C, D) or none (E), all of which were set to 1. Data are presented as means ± s.e.ms.; n = 3 [A (for 2xGamillus, 2xRFP, albumin-565, ALT-488), B (for 2xGamillus), D (for RFP-GFP), E (for 2xGamillus, 2xRFP, Amyloid-β-488, transferrin-488)], n = 4 [A (for Amyloid-β-488)], n = 5 [D (for Amyloid-β-488), E (for albumin-565)], n=6 (C), n =15 [B (for Amyloid-β-488)].

---

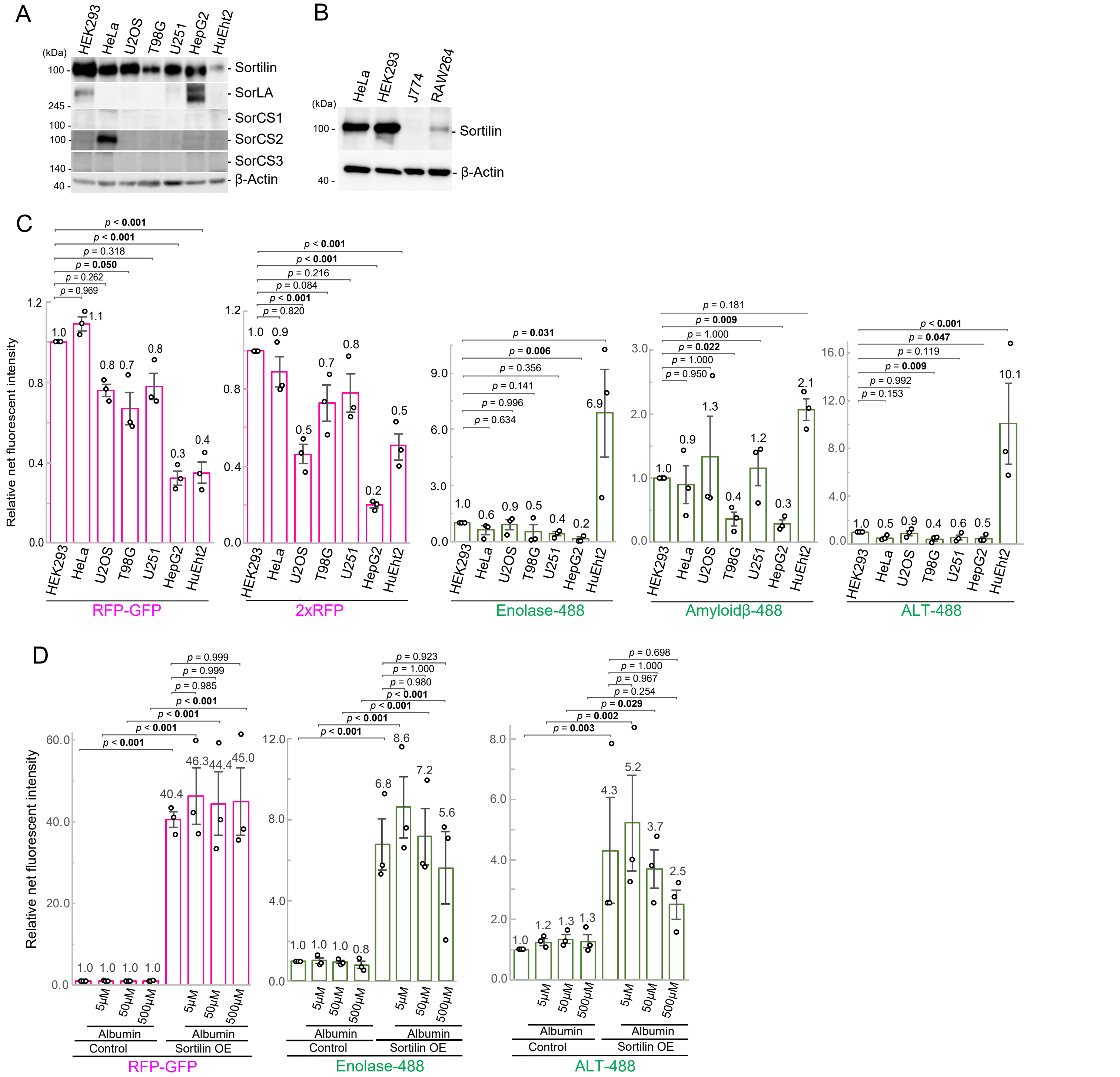


**Figure S3. Internalization of extracellular proteins in various cell lines.**
(A) Immunoblot of the endogenous sortilin family proteins in various cell lines using the indicated antibodies. (B) Immunoblot of endogenous sortilin in macrophage cell lines using the indicated antibodies. (C) FCM analysis of fluorescent extracellular protein internalization in various cell lines. The indicated cells were incubated with the indicated proteins for 16 h. Fluorescence intensity per cell was measured by FCM. (D) Competitive inhibition of extracellular protein internalization by albumin. Control or sortilin OE cells were incubated with indicated proteins in the presence or absence of 5-500 μM albumin for 16 h, and analyzed by FCM. The relative net fluorescent intensity is the -fold change of the net fluorescent intensity of each sample normalized to the net fluorescent intensity of HEK293 cells (C) or control cells (D), all of which were set to 1. Data are presented as means ± s.e.ms.; n = 3 (C, D).

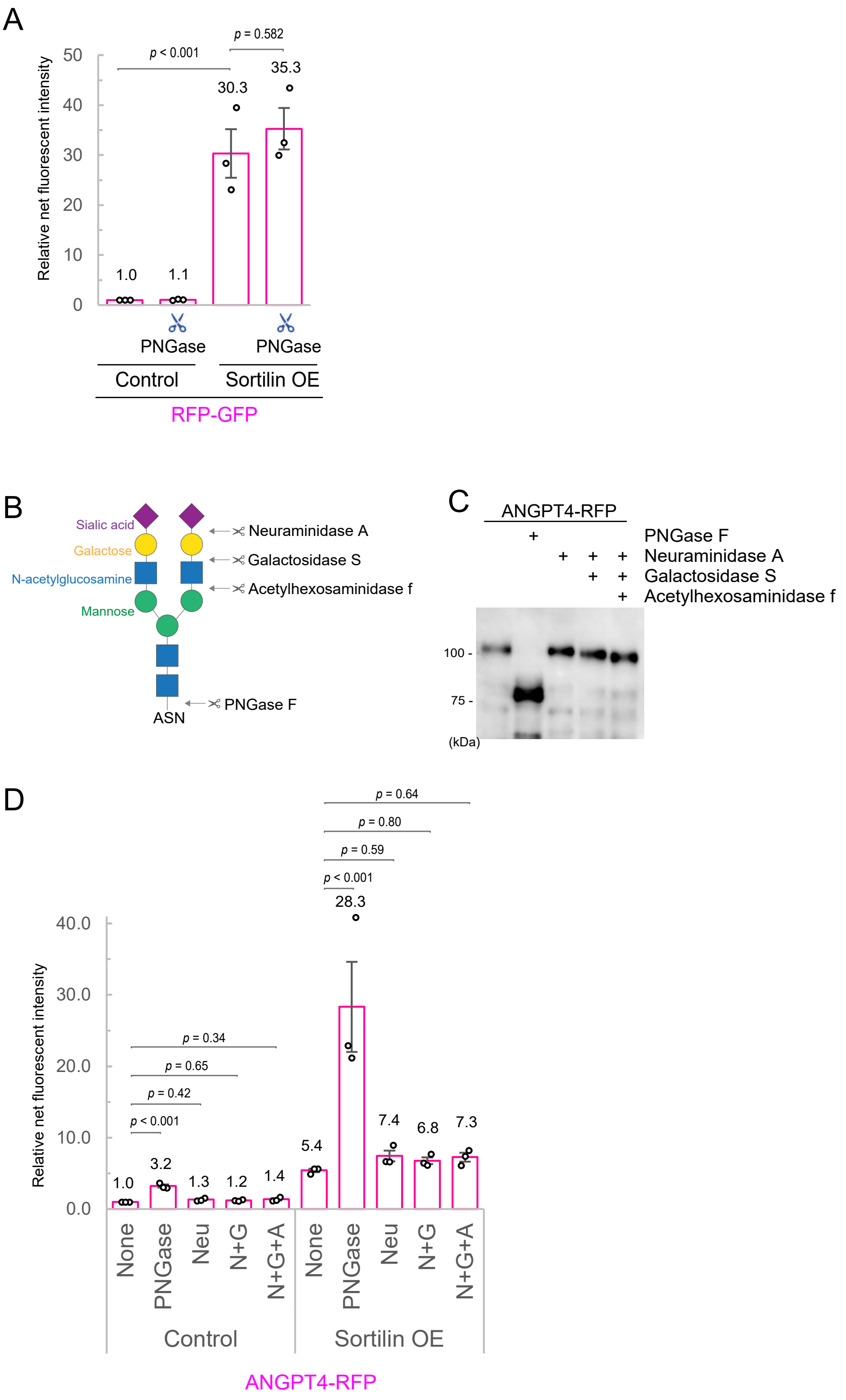


**Figure S4. Internalization is markedly enhanced by total glycan removal; partial glycan trimming was of little effect.**
(A) FCM analysis assessing the effect of PNGase F treatment on protein internalization. RFP-GFP was pre-treated with PNGase F for 3 h at 37°C. Control and sortilin OE cells (HEK293) were incubated with 0.5 µM RFP-GFP for 16 h, and analyzed by FCM. PNGase F treatment did not affect the internalization efficiency of RFP-GFP, indicating that the enzyme per se did not non-specifically affect sortilin-mediated internalization. (B) Schematic of the glycosidases. Neuraminidase A, galactosidase S, and acetylhexosaminidase f are exoglycosidases that specifically cleave terminal sialic acid, galactose, and N-acetylglucosamine residues respectively. In contrast, PNGase F is a glycosylamide hydrolase that cleaves N-glycans from polypeptide chains at the glycan roots. (C) Immunoblot of glycosidase-treated ANGPT4-RFP preparations. ANGPT4-RFP was treated with the indicated glycosidases for 3 h at 37°C and then subjected to immunoblotting using the anti-mCherry antibody. (D) FCM analysis of deglycosylated ANGPT4 internalization. ANGPT4-RFP was pre-treated with the indicated glycosidases for 3 h at 37°C. Control and sortilin OE cells (HEK293) were incubated with 0.2 µM ANGPT4-RFP for 16 h and analyzed by FCM. The relative fluorescent intensity is the -fold change of the net fluorescent intensity of each sample normalized to the net fluorescent intensity of control cells (A, D), both of which were set to 1. Data are presented as means ± s.e.ms.; n = 3 (A, D).

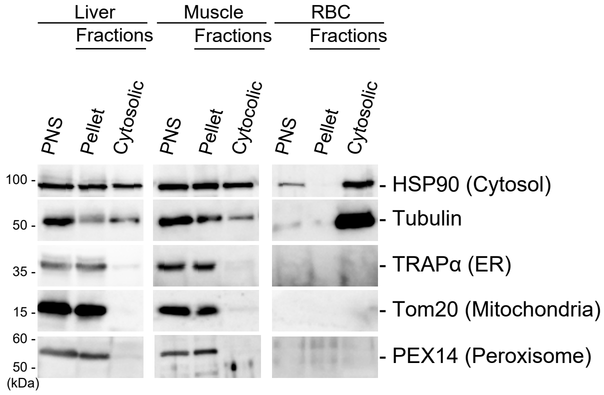


**Figure S5. Validation of subcellular fractionation to isolate the cytosolic fraction from mouse tissues.** Post-nuclear supernatants (PNS) from mouse liver, skeletal muscle, and red blood cells (RBC) were fractionated into membrane/organellar pellets (Pellet) and cytosolic fractions (Cytosolic) by ultracentrifugation. Fractions were analyzed by immunoblotting using markers for the cytosol (HSP90, Tubulin), endoplasmic reticulum (TRAPα), mitochondria (Tom20), and peroxisome (PEX14) to confirm the successful isolation of cytosolic proteins without organellar contamination.

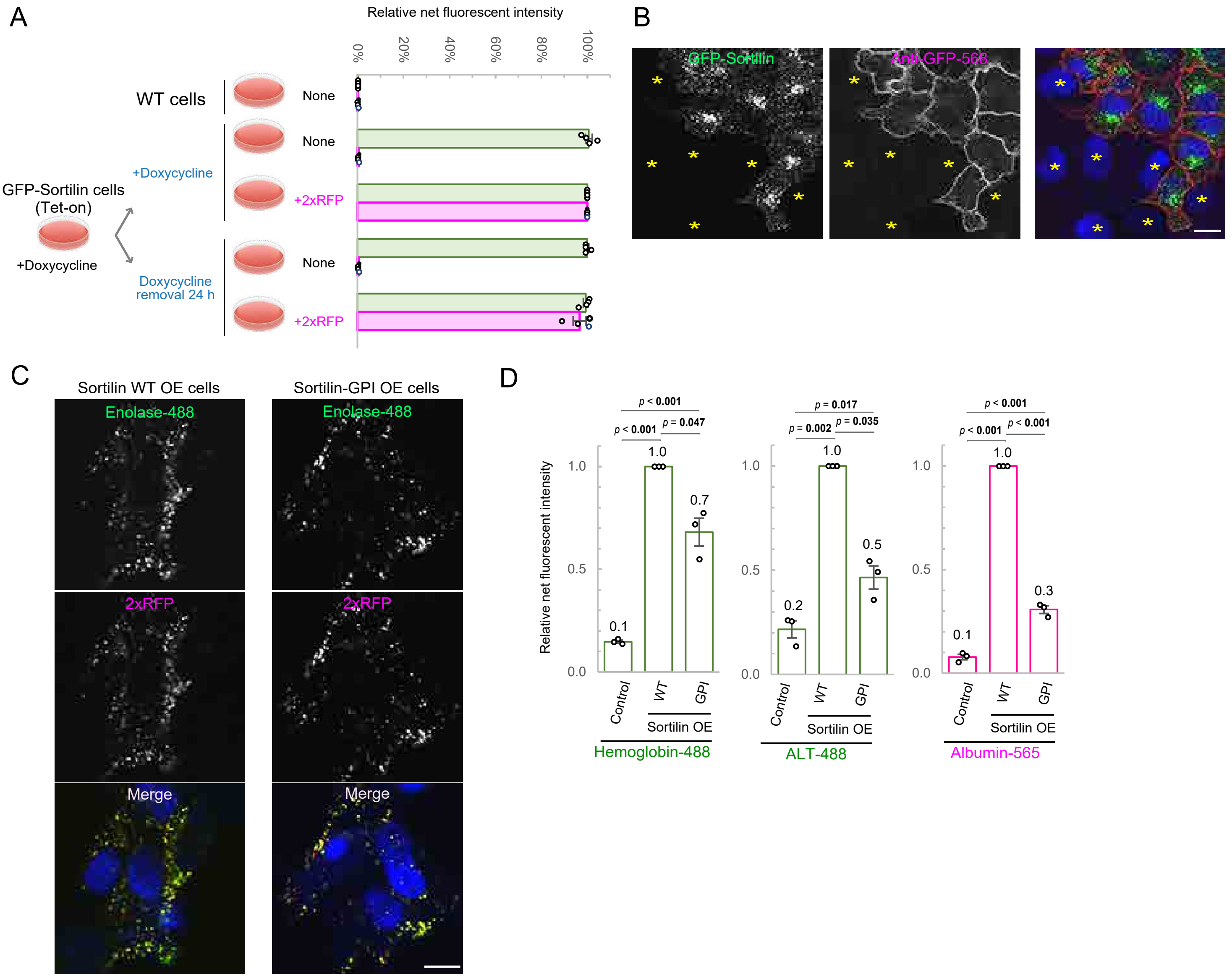


**Figure S6. Sortilin is not degraded during substrate internalization.**
(A) A schematic of sortilin stability analysis using the Tet-On system. sortilin-GFP (Tet-On) cells were initially treated with doxycycline (Dox) for several days to induce expression. Then, the cells were incubated for an additional 24 h under each of four conditions: with or without Dox, and in the presence or absence of 1 µM 2xRFP for 16 h. Right panel: The sustained receptor expression observed after Dox removal, regardless of the presence or absence of 2xRFP internalization, suggests that the receptor is not degraded with its substrate. Rather, the receptor likely undergoes efficient recycling after mediating substrate binding and subsequent lysosomal delivery. (B) Specific cell-surface labeling of GFP-sortilin at time 0. Confocal microscopy of GFP-sortilin–expressing HeLa cells immediately after surface labeling with Anti-GFP-568 at 4°C (time 0). Anti-GFP-568 signals are exclusively localized to the plasma membrane of GFP-sortilin–expressing cells and are completely absent in non-expressing control cells (asterisks). (C) Internalization of substrates in sortilin-GPI OE cells. Sortilin WT or -GPI OE cells (HEK293) were incubated with enolase-488 and 2xRFP for 16 h, fixed, and analyzed by confocal microscopy. Both enolase-488 and 2xRFP were detected in the intracellular region but not on the cell surface in both cell lines. Fluorescence detection sensitivity was not kept constant across images, as they were intended simply to show subcellular localization. (D) FCM analysis of substrate endocytosis as in Figure 7H with various substrates. Control cells and the indicated Sortilin OE cells (HEK293) were incubated with the indicated proteins for 16 h, and analyzed by FCM. The relative net fluorescent intensity in FCM is the -fold change in the net fluorescent intensity of each sample normalized to the net fluorescent intensity of control cells, all of which were set to 1. Scale bars: 10 µm.
